# Understory light quality affects leaf pigments and leaf phenology in different plant functional types

**DOI:** 10.1101/829036

**Authors:** CC Brelsford, M Trasser, T Paris, SM Hartikainen, TM Robson

**Affiliations:** Organismal and Evolutionary Biology, Viikki Plant Science Centre, Faculty of Biological and Environmental Sciences, University of Helsinki, Helsinki, Finland; Gregor Mendel Institute of Molecular Plant Biology, Vienna, Austria; Normandie Université, UNIROUEN, Ecodiv, Rouen, France

**Keywords:** phenology, pigments, flavonoids, spectral composition, canopy shade

## Abstract

Understory plant species take on different functional strategies, whereby some exploit periods of available light in springtime before the canopy closes, and others also benefit from sunlight later in autumn when the canopy opens again. These strategies involve understory species coordinating phenological events to pre-empt canopy leaf out and to extend their growing season beyond canopy leaf senescence, meanwhile accumulating photo-protective pigments which mitigate periods of high-light exposure. Canopy closure brings shade to the understory, but also causes drastic changes in light quality. Whilst many experiments manipulating spectral quality have revealed understory plant responses to the changing R:FR ratio in shade, effect of the blue and UV regions have been examined very little. We installed filters attenuating short wavelength regions of the solar spectrum in a forest understory in southern Finland, creating the following treatments: a transparent control filter, and filters attenuating UV radiation < 350 nm, all UV radiation, and both UV and blue light. In eight understory species, representing different plant functional types, we repeatedly assessed leaf optical properties to obtain epidermal flavonol and anthocyanin contents from leaf emergence in spring to leaf senescence in autumn, during both 2017 and 2018. Flavonols responded more to seasonal changes in light quality in relatively light-demanding species than in shade-tolerant and wintergreen species; and were particularly responsive to blue light. However, anthocyanins were largely unaffected by our filter treatments, suggesting that other cues such as cold temperatures govern their seasonal variation. UV radiation only accelerated leaf senescence in *Acer platanoides* seedlings, but blue light accelerated leaf senescence in all species measured apart from *Quercus robur*. In summary, seasonal changes in understory solar radiation in the blue and UV regions affected leaf pigments and leaf phenology; particularly for more light-demanding species. An increase in canopy duration under climate change will extend the period of shade in the understory, with consequences for the spectral cues available to understory plants. The resultant reduction in blue and UV radiation in shade, could delay leaf senescence in the understory even further.

## Introduction

Forest understories are dynamic and heterogeneous light environments. Many plant species are able exploit increasing sunlight reaching the understory prior to canopy closure for photosynthesis during early spring, while some also extend their growing season to exploit favorable conditions after canopy leaf senescence in the autumn (Kudo et al. 2008, Richardson and O’Keefe 2009). Different plant species segregate into functional types with differing light-capture strategies within the forest understory plant community (Grubb 1977, Wolkovich and Cleland 2011, Heberling et al. 2019). Accordingly, species may adopt different strategies to time their growth and life cycle, adapting to seasonal variation in light availability in the forest understory (Augspurger et al. 2005, Heberling et al. 2019). These seasonal adaptations partially lead different species to grow beneath deciduous and evergreen canopies (Frelich et al. 2003).

Environmental cues help plants time their phenology, (Chuine and Régnière 2017, Flynn and Wolkovich, 2018). Spring ephemeral species time their emergence and leaf out in spring to maximize carbon gain then senesce during canopy closure when the irradiance drops in deciduous stands (Kudo et al. 2008). During autumn leaf senescence, the canopy opens, and once again more light is available to the forest understory (Richardson and O’Keefe 2009). This autumn light niche in the understory is exploited by many perennial herbaceous species and tree seedlings which delay leaf senescence until during or after canopy opening, extending photosynthesis for overwinter carbon storage (Augspurger et al. 2005, Kudo et al. 2008). Wintergreen plants produce new leaves each year which overwinter then senesce during the growing season (Heberling et al. 2019). Although carbon gain is often negligible during winter, this strategy allows them to exploit favorable conditions for photosynthesis in early spring during snowmelt (Landhäusser et al. 1997, Saarinen et al. 2015). Shade-tolerant species in the understory may likewise have later leaf emergence, but extend their leaf longevity utilizing low light efficiently, whereas the leaves of more light-demanding species may emerge earlier to exploit relatively high understory irradiances before canopy closure (Niinemets, 2010).

Once the canopy closes, the understory light environment is mostly shade (Chazdon and Pearcy, 1991). Not only does the amount of irradiance change in the understory, but so does its spectral composition, the light quality (Federer and Taner 1966, Vezina et al. 1966, Ross and Flanagan 1986, Messier et al. 1998, Leuchner et al. 2005, Navrátil et al.2007, Hartikainen et al. 2018). Far-Red (FR) light is mostly reflected or transmitted from canopy leaf surfaces and scattered into the understory shade, reducing the ratio of red(R): FR light (Ballaré and Pierik 2017). These changes in R:FR in the understory are detected by phytochrome photoreceptors, which coordinate the shade avoidance syndrome in many species (Ballaré et al. 1987). Similarly, the irradiance of blue light and UV radiation penetrating into the forest understory is reduced during canopy leaf out, due to its absorption by the plant canopy (Casal 2012), although, in relative terms, the ratio of UV: photosynthetically active radiation (PAR) is increased in understory shade due to the higher diffusivity of UV radiation (Flint and Caldwell 1998, Grant et al. 2005).

Plants produce photo-protective pigments such as carotenoids and flavonoids in response to highlight stress and UV radiation (Agati and Tattini 2010, Agati et al. 2012, Brunetti et al. 2018). Flavonoid accumulation in response to blue light and UV-A is governed by CRYs, but UVR8 dictates plant response to UV-B radiation (Wade et al. 2003). The action spectra of these two photoreceptors may in fact overlap in the UV-A region (Morales et al. 2013, Brelsford et al. 2018). Rai et al. (2019) conjecture that responses, like flavonoid induction, primarily governed by CRY or UVR8 segregate at 350 nm wavelength. Flavonoids can be broadly separated into anthocyanins and flavonols/flavones (Agati et al. 2012); both classes function as antioxidants with epidermal flavonols also screening UV-radiation, although anthocyanins also absorb in the blue and to a lesser extent UV region of the spectrum (Steyn et al. 2002).

We know relatively little about how the role of flavonoids differs among plant functional types in response to light. It has been suggested that epidermal UV-screening is highest in evergreen plants, intermediate in deciduous woody plants, and lowest in herbaceous plants (Day 1992, Day 1993, Semerdjieva et al. 2003, Li et al. 2010). However, given that relatively few studies have been designed explicitly to compare flavonoid accumulation among different plant functional types, we lack the information needed to make these broad generalizations. There are several hypotheses identifying different environmental cues accounting for the seasonal dynamics of anthocyanins, including temperature, herbivory and light stress (Lee 2002, Hoch et al. 2003, Karageorgou and Manetas 2006, Archetti et al. 2009). For many plant species, anthocyanins increase in senescing leaves (Feild et al. 2001, Lee 2002), and shading can reduce the production of anthocyanins during autumn in woody species, as well as delaying leaf senescence, irrespective of the R:FR ratio (Lee et al. 2003). To date, it is still unknown what light signal, if any, is responsible for the reduction of autumnal anthocyanins and delayed leaf senescence under shaded conditions. Interestingly, supplementary UV radiation from lamps has been shown to accelerate leaf senescence in *Fagus sylvatica*, possibly through increased photodamage throughout the growing season (Zeuthen et al. 1997). As such, the reduction of UV radiation under canopy shade could affect leaf senescence in understorey plant species. Furthermore, climate change is expected to increase canopy duration, with earlier canopy leaf out and later leaf senescence (Piao et al. 2019, Vitasse et al. 2011, Buitenwerf et al. 2015). However, it is not clear how the changes in light quality under a prolonged period of canopy shading, will affect leaf pigments and the leaf phenology of understory plants.

Here, we examine how depleted solar blue light and UV radiation affect the seasonal dynamics of leaf pigments and phenology, by following different understory plant species over a 2-year period. We tested four hypotheses:

1. Based on our knowledge of flavonoid responses to light quality, attenuating blue light should cause the greatest reduction in flavonoid accumulation.
2. Flavonoid accumulation should be more responsive to blue and UV attenuation treatments, in relatively light-demanding species than in more shade-tolerant species. We expect this because more light-demanding species need to exploit periods of seasonally high irradiance, so require greater plasticity to readily adjust their photoprotection.
3. Changes in light quality during spring would have a greater effect on phenology of tree seedlings than deciduous herbaceous species in the understory. This is because the buds of tree seedlings are exposed to the sunlight above-ground during winter and spring when deciduous herbaceous species are largely dormant below-ground.
4. Attenuating UV radiation would reduce photo-damage to leaves through the season and thus delay autumn leaf senescence.

## Methods

### Climate and site information

Our experiment was conducted at Lammi Biological Station, situated at 61.05°N, 25.05°E. The average monthly temperature and annual precipitation were 5.1°C and 572.4 mm respectively between 2017 and 2018. Daily mean, minimum and maximum temperatures (Fig. S1), precipitation (Fig. S2), and solar radiation (Fig. S3, Fig. S4) were recorded at the site and processed by the Finnish Meteorological Institute (https://en.ilmatieteenlaitos.fi/weather/h%C3%A4meenlinna/lammi).

The effect of canopy closure on understory spectral composition in the stands was measured in 2015 (Hartikainen et al. 2018), and during canopy leaf-out between April and June in 2016, 2017, and 2018. Spectral irradiance was recorded with a Maya 2000 Pro spectrometer (Ocean Optics, Dunedin, FL, USA) calibrated for accuracy in the solar UV and PAR regions of the spectrum. The same device and protocol as Hartikainen et al. (2018) were used for measurements and processing of the irradiance data. Ambient solar PAR above the canopy at Lammi Biological Station between 2015-2019 was also recorded to show the effects of changing cloudiness on incoming irradiance (PQS1 PAR Quantum Sensor, Kipp & Zonen, Delft, Netherlands, Fig. S4).

### Experimental design

Our experimental plots were installed under two stand types, deciduous stands dominated by either *Quercus robur* or *Betula pendula*, and an evergreen stand of *Picea abies*. We used 15 split-plots; six under *P. abies* (hereafter referred to as the evergreen stand) and nine in the deciduous stands. Our plots were adjacent to the plots used by Hartkainen et al. (2018), with the same canopy species, stand spacing, architecture and LAI (Table S2).

One replicate from each of four different filter treatments, made from polycarbonate (Foiltek Oy, Vantaa, Finland), was randomly arranged within every plot. The control filter was equally transparent to all wavelengths of solar radiation (PLEXIGLAS 2458 GT SOLARIUMLAATU), one filter treatment attenuated UV radiation below 350 nm (PLEXIGLAS 0Z023), one filter treatment attenuated all UV radiation (ARLA MAKROLIFE), and one filter treatment attenuated blue light and all UV radiation (PLEXIGLAS 1C33 (303)). The filter dimensions were 88 cm length x 60 cm width x 40 cm in height. All filters were orientated with the longest-sloping side facing south, and short vertical side facing north, with an air vent at the apex to increase air flow and reduce warming (Fig. S5). Each filter was also raised on 10-cm wooden blocks to allow airflow. We allowed a 10-cm border around a central area inside the filters where plants were measured, to avoid filter-edge effects. (Aphalo et al. 2012).

### Selection of understory species

To capture responses from different plant functional types in the forest understory, we used volunteer plant species already growing beneath the installed filters (Table 1): *Aegopodium podagraria, Anemone nemorosa, Fragaria vesca, Mainthemum bifolium, Oxalis acetosella* and *Ranunculus cassubicus. R. cassubicus, A. podagararia* and *Q. robur* were only present in deciduous stands, and *M. bifolium* was only present in the evergreen stand. We also transplanted tree seedlings from two target species, *A. platanoides* and *Q. robur* within the same stand. Germinating seedlings were transplanted at the two-cotyledon stage, *A. platanoides* in April 2016 and *Q. robur* in September 2017. Plants of similar size were transplanted into areas under the filters with the least existing vegetation to avoid disturbance. Four *A. platanoides* seedlings were transplanted under each filter, and four *Q. robur* seedlings were transplanted under each filter in the deciduous stands. *Q. robur* seedlings were not present in the evergreen stand and so were not transplanted there.

**Table 1.** Summary of data collected from understory species during the experiment, their functional strategies, and the number of filters that they were present under.

| Species | Stand types | Pigment measurements | Phenology measurements | Plant Functional Type | Light strategy | Phenological strategy | Present under ## filters |
| --- | --- | --- | --- | --- | --- | --- | --- |
| *A. nemorosa* | Deciduous and evergreen | 2017 | Spring 2018 | Herbaceous | More light-demanding | Spring ephemeral | 49 |
| *A. platanoides* | Deciduous and evergreen | 2017 and 2018 | Spring and autumn 2017 and 2018 | Tree seedling | More light-demanding | Autumn senescing | 59 |
| *A. podagraria* | Deciduous | 2017 and 2018 | Spring and autumn 2018 | Herbaceous | More light-demanding | Autumn senescing | 25 |
| *F. vesca* | Deciduous and evergreen | 2017 and 2018 | - | Herbaceous | Shade tolerant | Wintergreen/heterotypic | 52 |
| *M. bifolium* | Deciduous and evergreen | 2017 and 2018 | - | Herbaceous | Shade tolerant | Summer green | 22 |
| *O. acetosella* | Deciduous and evergreen | 2017 and 2018 | - | Herbaceous | Shade tolerant | Wintergreen | 41 |
| *Q. robur* | Deciduous | 2018 | Autumn 2018 | Tree seedling | More light-demanding | Autumn senescing | 23 |
| *R. cassubicus* | Deciduous | 2018 | Spring and autumn 2018 | Herbaceous | More light-demanding | Autumn senescing | 20 |

The filters partially blocked precipitation from reaching the ground underneath them, so additional watering was provided to the plants every 3 days (Fig. S6). The soil moisture in the plot was also monitored with a TDR probe (SM200 Moisture Sensor with HH2 Moisture Meter, Delta-T Devices, Cambridge, UK), and we ensured soil moisture was equivalent amongst treatments (Fig. S6). Air temperature, monitored with iButton sensors (Maxim Integrated, San Jose, United States), was on average 0.3 °C higher under the filters than in the ambient understory (Fig. S7), with no difference between treatments or filter types (Fig. S7).

### Measurements of leaf pigments and leaf phenology

A Dualex Scientific+ (Force-A, University Paris-Sud, Orsy, France) was used to make nondestructive measurements of leaf pigments (chlorophyll content, and epidermal flavonols and anothcyanins) based on their optical properties (Cerovic et al., 2012). Both adaxial and abaxial measurements of flavonol and anthocyanins were made every week between May 15^th^ and Oct 2nd 2017, and on five occasions between May 14th and October 1^st^ in 2018. For plants with few leaves, namely *A. platanoides, A. nemorosa, Q. robur, M.bifolium*, pigments were measured from all the leaves on each plant. For *A. podgraria, F. vesca* and *O. acetosella* two leaves per plant were measured. After the production of new leaves in spring, the same cohort of leaves were followed until the last measuring dates in October 2017 and 2018 respectively, at which point the proportion of leaves which had senesced was too high for continued measurement.

Bud burst and leaf out of *A. platanoides* was measured every 2-3 days, along a scale of 1-7 adapted from Teissier du Crois et al. (1981); whereby 1 = dormant, 2 = bud swelling, 3 = bud split, 4= leaf tip protruding, 5=leaf mostly out, 6= leaf out but not fully expanded, and 7 = fully expanded leaves. For the herbaceous species, *A. podagraria, A. nemorosa, R. cassubicus*, emergence of new leaves was measured once a week as 1 = shoot visible, 2= expanded leaf. Leaf senescence for all tree and herbaceous species was measured on a scale of 1-5, whereby 1=fully green, 2=starting to yellow, 3=mostly yellow, 4=turning brown, 5=all leaves fallen. Leaf senescence was censured for all the leaves on each plant every 2 weeks. Bud burst was not recorded in *Q. robur*, which only germinated in spring 2018. Canopy opening and canopy closure were recorded in the deciduous stands in both years, and determined as the date when the canopy had reached 50% leaf out (canopy closure) and 50% leaf fall (canopy opening).

To determine whether understory spectral composition affected the maximum quantum yield of photosystem II (PSII) photochemistry (*F*_v_/*F*_m_), leaves of *A. platanoides* were dark-adapted for 30 min using darkening leaf clips and measured with a mini-PAM fluorometer (Heinz-Walz GmbH, Effeltrich, Germany) on the 24^th^ −25th June 2017 between 10:00-16:00. Stomatal conductance (*g*_s_) was measured in *A. platanoides* using an AP4 leaf porometer (Delta-T Devices, Cambridge, UK), between 10:00-14:00 under clear sky conditions on the 16th June 2018. In both cases, two leaves were measured per plant. A summary of all measurements made is given in Table 1 – time constraints limited pigment and phenological measurements of some species to just one year and physiological measurements to just one time point.

#### Statistical methods

For all statistics, the unit of replication was at the level of the filter treatment per plot. Model selection was based on meeting the requirements for statistical analyses laid out by Zuur et al. (2010). An LME model was used to test the effects of Treatment x Stand type x Time as fixed factors, and with a nested random effects structure of Time/Stand type/Stand/Plot ID, using the package ‘nlme’ in R Studio (Pinheiro et al. 2017). When there were non-linear trends over time, producing heteroscedasticity in residuals, a mixed effects generalized additive model (GAMM) (package = ‘mgcv’, Wood and Wood 2015), was used with Treatment x Stand Type as parametric terms, and a smoothing term for time, as well as a smoothing term for the random effects structure (Time/Stand type/Stand/Plot ID) using the function bs=‘re’ as described by (Pedersen et al. 2019). The type of spline used as the smoothing term for time was chosen based upon the model which had the lowest AIC value. If heteroscedasticity in the residuals was present due to heterogeneous variation, then the weighting function ‘weights = varPower’ was used. If filter treatment was significant in the model, then the effect of the filter attenuating both blue light and UV was compared against the filter attenuating UV, to determine the effect of blue light. Similarly, the effect of the filter treatment attenuating UV was compared against the filter treatment attenuating UV below 350nm, to determine the effect of UV radiation above 350nm. The effect of UV radiation below 350nm was determined by comparing the filter attenuating UV below 350nm against the control treatment. These contrasts cannot distinguish whether combining different regions of the spectrum had additive or synergistic effects. When multiple tests were used, *p* values were corrected using Holm’s correction. A factor or an interaction between factors was considered significant in the model when *p*<0.05.

## Results

### Attenuating blue light delays leaf out in *A. platanoides*, but not in herbaceous species

There was a significant effect of filter treatment on leaf out of *A. platanoides* seedlings (Table S3 and S4). Attenuating blue light significantly delayed leaf out, whilst both attenuating UV radiation above and below 350nm wavelength had no significant effect. Averaged across both years, the date of bud burst was delayed by 1.4 days when blue light was attenuated, and final leaf out was delayed by 2.8 days when blue light was attenuated. Stand type also had a significant effect on phenology, as bud burst of *A. platanoides* was 5 days later in the evergreen stand than the deciduous stands. Filter treatments had no significant effect on the leaf out of any herbaceous species (Tables S5-S8), although stand type had a significant effect on *A. nemorosa* phenology, whereby its emergence (stage 1) occurred 7 days later in the evergreen stand compared to the deciduous stand (Table S5 and S6).

### Attenuating UV and blue light delays leaf senescence

Overall, attenuation UV radiation below 350nm significantly delayed leaf senescence in *A. platanoides* seedlings (Tables S9, S10 Fig 4). The onset of leaf senescence (stage 2) was delayed by 4.1 days when UV radiation below 350nm was attenuated, although final leaf fall (senescence stage 5) occurred on the same day irrespective of filter treatment (Fig 4). In comparison, attenuating blue light and UV radiation above 350nm had no overall effect on leaf senescence in *A. platanoides* seedlings (Table S10). However, in the evergreen stand attenuating blue light delayed the onset of leaf senescence in *A. platanoides* seedlings by 14.3 days, and final leaf senescence by 7.6 days. Attenuation of blue light also delayed the onset of leaf senescence in herbaceous species *R. cassubicus* by 7.4 days, and final leaf senescence by 7.0 days (Tables S11, S12). There was no significant effect of UV radiation below or above 350nm on leaf senescence in *A. podagraria* (Table S13, S14). This was because, although *A. podagraria* growing under the filter treatment attenuating blue light and UV had significantly later leaf senescence than under the control treatment, the effect of the blue light was not significantly different from the no UV treatment (Table S13, S14). This suggests that blue light and UV radiation may have an additive effect on leaf senescence in *A. podagraria*. The onset of leaf senescence in *A. podagraria* was delayed by 5.6 days beneath the filter attenuating blue light and UV radiation compared to the control treatment, however final leaf senescence occurred on the same day. Our filter treatments had no effect on leaf senescence in *Q. robur* seedlings (Table S15, Fig. 3).

**Figure 1.**
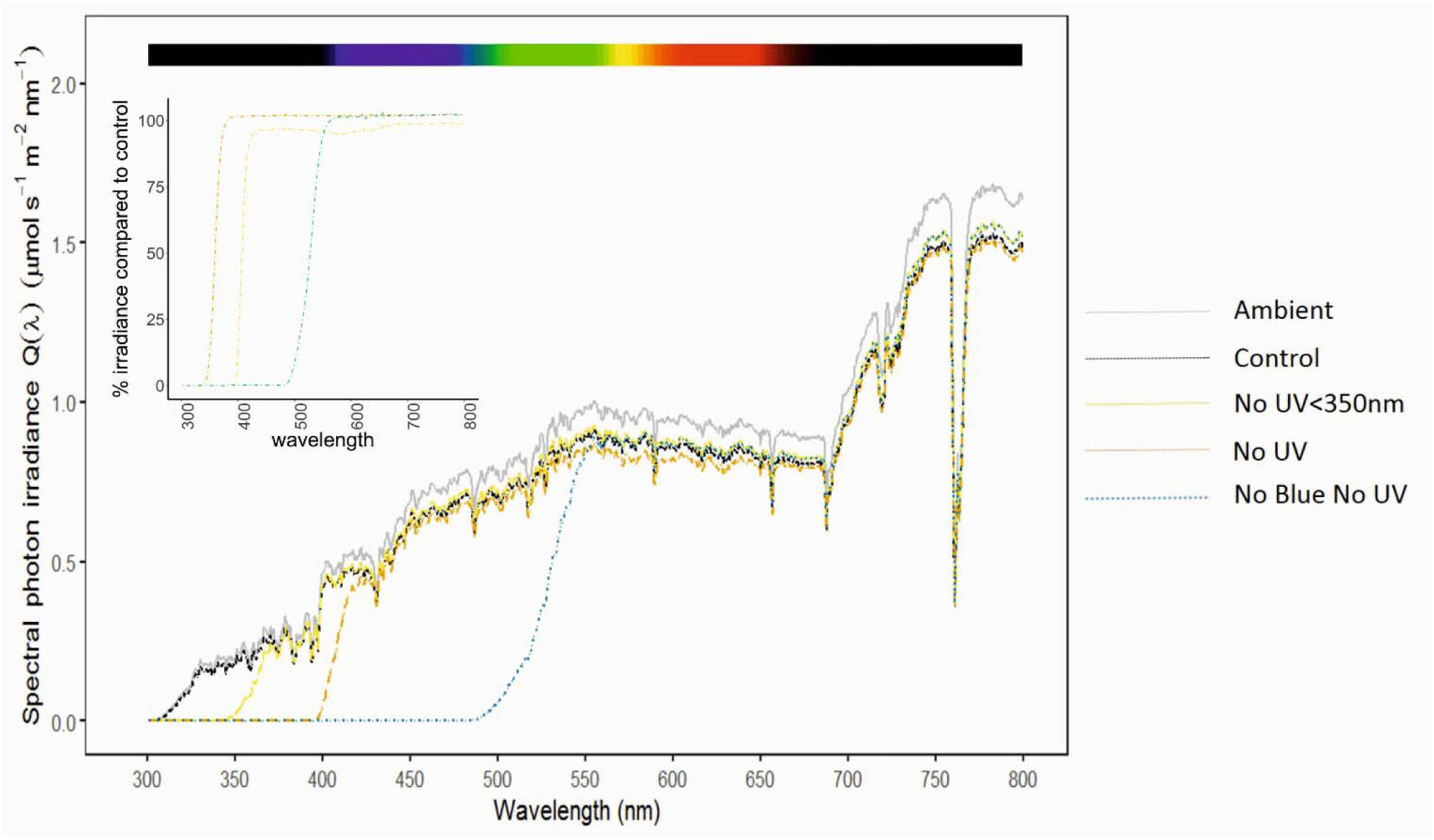
Understory spectral irradiance in canopy shade during spring leaf out (25/04/2015). The measurements were taken in a deciduous *B. pendula* stand, following methods described in Hartikainen et al. (2018). The different filter treatments are represented by line colours & types, whereby grey = ambient understory irradiance, black = transparent control filter, yellow = attenuation of UV radiation below 350nm, orange = attenuation of total UV radiation, and blue= attenuation of blue and UV radiation. Spectral irradiance measurements under the forest stands between 2015 and 2018 are shown in supplemental Table S1.

**Figure 2.**
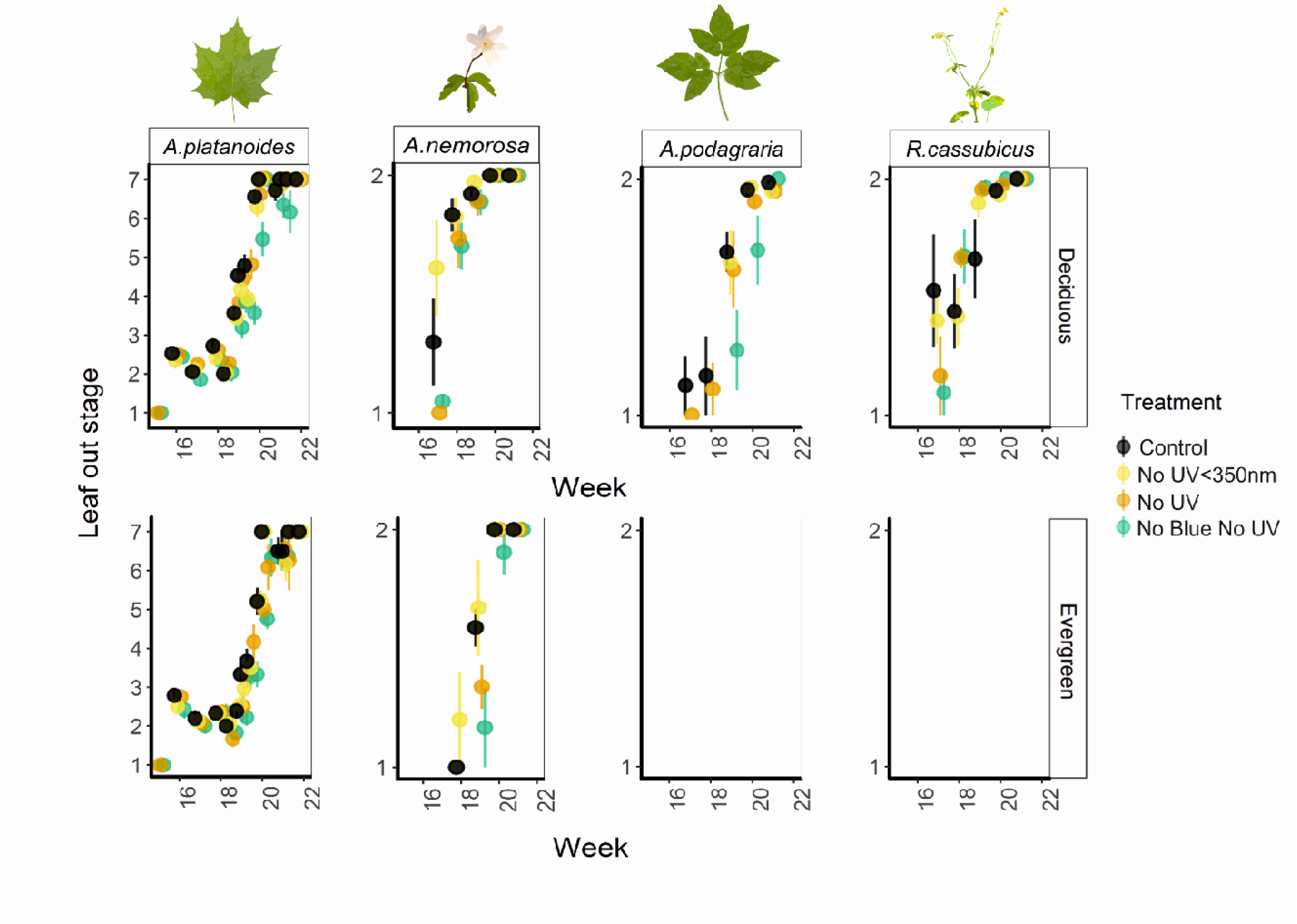
Spring leaf out of understory plant species growing beneath filters attenuating different regions of the solar spectrum. For *A. platanoides* leaf out was scored on a scale of 1-7 whereby 1 = dormant, 2 = bud swelling, 3 = bud split, 4= leaf tip protruding, 5=leaf mostly out, 6= leaf out but not fully expanded, and 7 = fully expanded leaves. For the other species, leaf out was scored on a scale of 1-2 whereby 1 = shoot visible, 2= expanded leaf.

**Figure 3.**
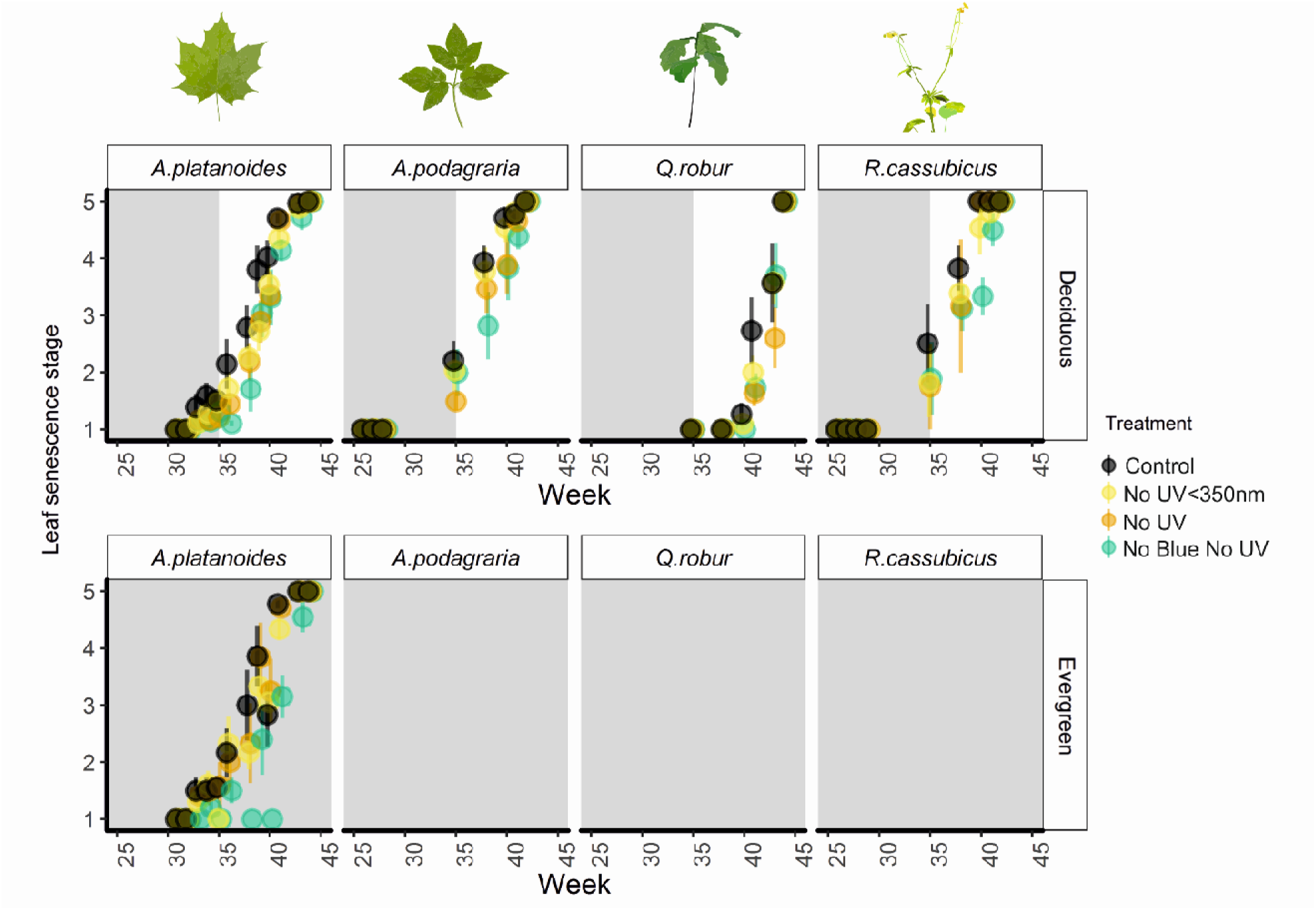
Leaf senescence stage for five understory plant species growing during 2017 and 2018. Leaf senescence was scored on a scale of 1-5, whereby, 1=green leaf, 2=starting to yellow, 3=mostly yellow, 4=mostly brown, 5=leaf fall. The grey shaded area represents the time that canopy was closed during the measurement period. Means and ± 1 SE presented on the graph, with the individual plot as the unit of replication. Where species were absent from the evergreen stand panels were left blank.

**Figure 4.**
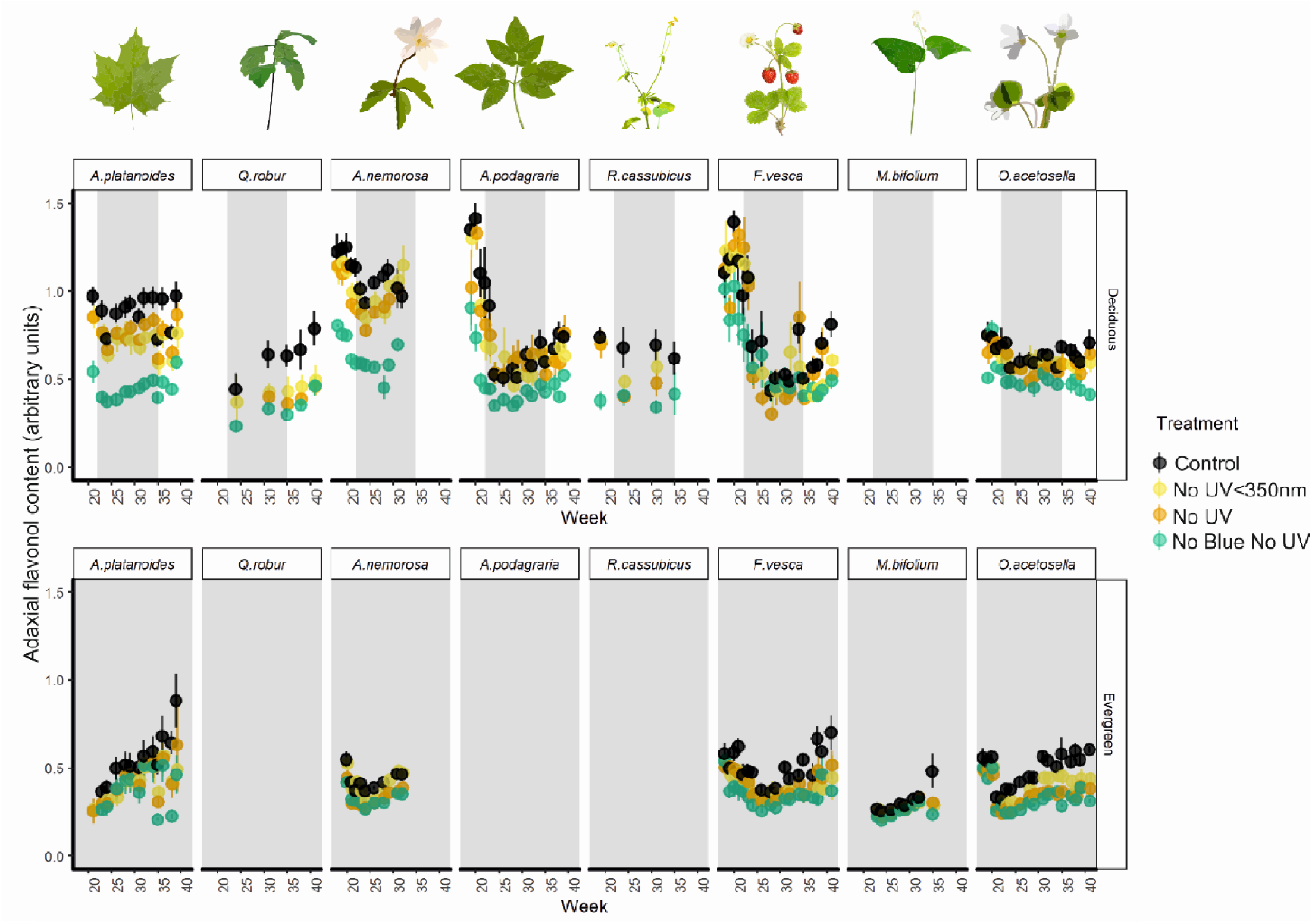
Adaxial epidermal flavonol content of eight different understory plant species, growing in deciduous and evergreen stands during 2017 and 2018. The grey shaded area represents the period that canopy was closed during the measurement period. Means and 1 ± SE are presented on the graph, with the individual filter as the unit of replication. Where species were absent from the stands panels were left blank.

### Attenuating blue light reduces flavonol accumulation the most in light-demanding species

For most understory plant species we monitored in the deciduous stands, the general trend was for high adaxial epidermal flavonol content during spring, followed by a decrease during the period of canopy closure, before it increased again during autumn leaf senescence (Fig 5). Despite the absence of canopy closure and opening, a similar but less pronounced seasonal pattern in adaxial epidermal flavonol content was also present in understory species growing in the evergreen stands. Likewise, while our different filter treatments reduced flavonol content to differing extents, the overall seasonal trends remained consistent among all filter treatments.

**Figure 5.**
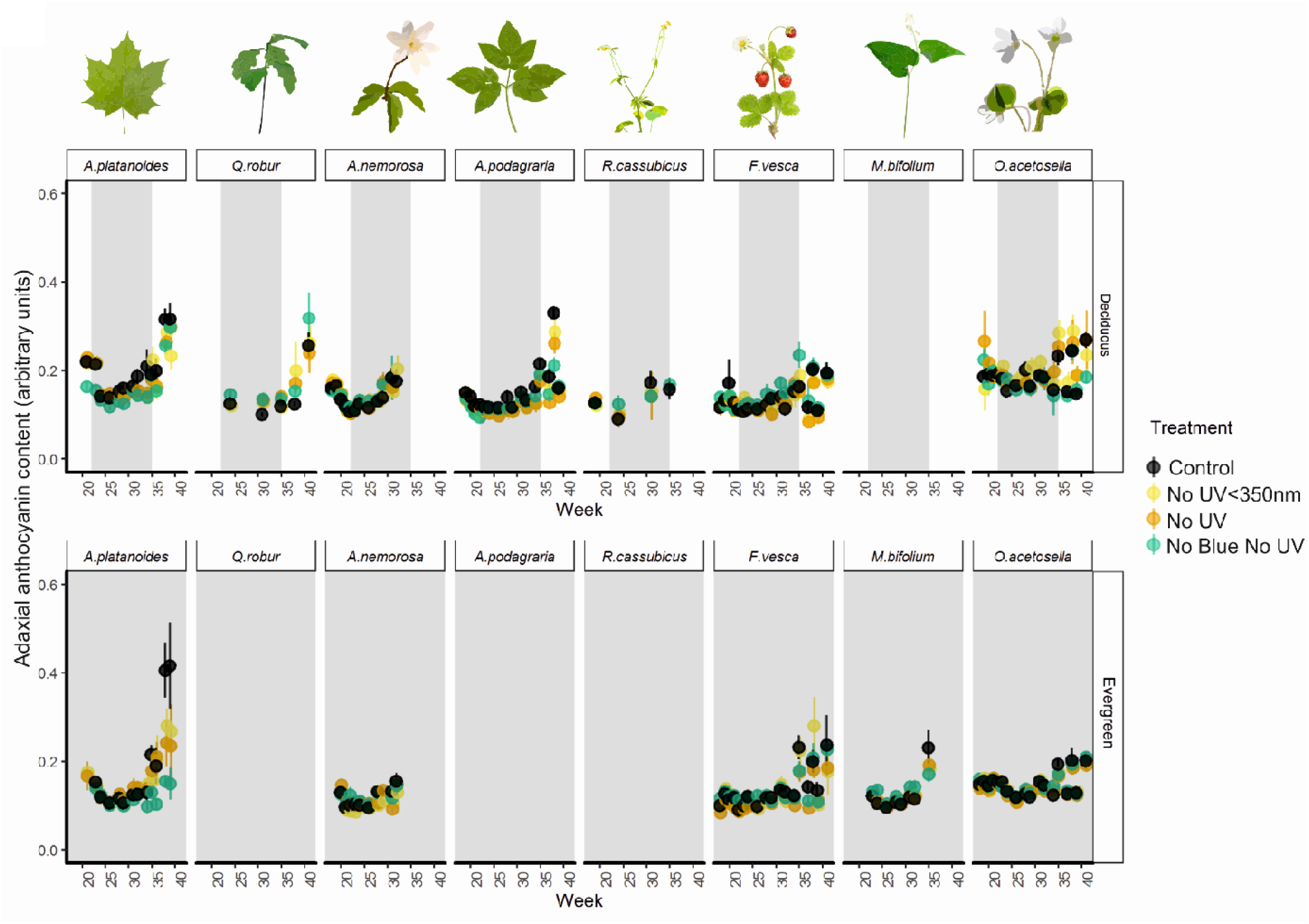
Adaxial epidermal anthocyanin content of eight different understory plant species, in our understory filter treatments. The grey shaded area represents the period during which the canopy was closed. Means and 1 ± SE are presented on the graph, with the individual filter as the unit of replication. Where species were absent from the stands panels were left blank.

Comparing the effects of spectral regions calculated from the differences between pairs of filter treatments, attenuation of blue light generally had the largest effect on adaxial epidermal flavonol accumulation (Table 2, Table S16A-P, Fig. 4). Attenuating blue light significantly reduced adaxial flavonols more than any other spectral region in the more light-demanding species, *A. nemorosa, A. platanoides, A. podagraria* and *R. cassubicus* (Table S16A-P, Fig. 4). Attenuating blue light also reduced adaxial flavonol accumulation in *M. bifolium* and *O. acetosella* (Table 2, Table S16G-L), although its effect was smaller in these shade-tolerant and wintergreen species (Table 2, Fig. 4).

**Table 2.** Effects of different regions of solar spectrum on average adaxial leaf pigment content of tree seedlings and herbaceous understory species throughout 2017 and 2018. The overall percentage change from plants receiving near-ambient solar radiation under the control treatments is given for both flavonols and anthocyanins. * = *p* <0.05, ** = *p* <0.01, *** = *p* <0.001, further details shown in Tables S16-S17.

| Species | Effect | Flavonol<br>difference | Flavonol<br>% | Sig | Anthocyanin<br>difference | Anthocyanin<br>% | Sig |
| --- | --- | --- | --- | --- | --- | --- | --- |
| *A. nemorosa* | blue | 0.41 | -25.63 | *** | -0.001 | -0.46 | ns |
|  | UV>350 | 0.09 | -5.43 | ** | -0.001 | 1.80 | ns |
|  | UV<350 | 0.08 | -10.12 | ns | 0.001 | -0.77 | ns |
| *A. platanoides* | blue | 0.32 | -31.03 | *** | 0.036 | -13.61 | * |
|  | UV>350 | 0.05 | 10.93 | *** | 0.008 | 3.32 | ns |
|  | UV<350 | 0.15 | -22.35 | *** | 0.016 | -8.97 | *** |
| *A. podagraria* | blue | 0.40 | -32.72 | *** | 0.021 | 0.18 | ns |
|  | UV>350 | 0.06 | 2.01 | ns | 0.018 | -4.19 | ns |
|  | UV<350 | 0.12 | -15.32 | ns | 0.013 | -9.87 | ** |
| *F. vesca* | blue | 0.16 | -3.93 | ns | 0.001 | 9.45 | ns |
|  | UV>350 | 0.10 | -8.37 | ns | 0.013 | -6.01 | ns |
|  | UV<350 | 0.06 | -10.64 | ** | 0.005 | -3.96 | ns |
| *M. bifolium* | blue | 0.05 | -6.11 | * | -0.011 | 7.57 | ** |
|  | UV>350 | 0.03 | 0.08 | ns | -0.002 | -0.85 | ns |
|  | UV<350 | 0.03 | -11.40 | ** | -0.003 | 2.57 | ns |
| *O. acetosella* | blue | 0.17 | -8.74 | *** | 0.003 | 1.19 | ** |
|  | UV>350 | 0.08 | -6.89 | ns | 0.004 | -3.29 | ns |
|  | UV<350 | 0.07 | -16.32 | ns | -0.001 | 0.46 | ** |
| *Q. robur* | blue | 0.33 | -6.88 | ns | -0.046 | 15.41 | ns |
|  | UV>350 | 0.15 | -4.13 | ns | -0.028 | -12.96 | ns |
|  | UV<350 | 0.15 | -36.01 | *** | -0.048 | 26.71 | ns |
| *R. cassubicus* | blue | 0.31 | -29.79 | * | 0.000 | 4.19 | ns |
|  | UV>350 | 0.06 | -3.65 | ns | 0.005 | 0.31 | ns |
|  | UV<350 | 0.07 | -11.52 | ns | 0.005 | -4.27 | ns |

Attenuating UV radiation below 350 nm significantly reduced adaxial epidermal flavonol content in *A nemorosa*, and surprisingly increased flavonol content *A. platanoides*, although both of these size effects were small (Table 2, Tables S16A-P). Attenuating UV radiation below 350 nm reduced adaxial epidermal flavonol content significantly in *A. platanoides, F. vesca, M. bifolium*, and *Q. robur* (Table 2, Tables S16A-P), and to a greater extent than longwave UV radiation (>350 nm) in these species (Table 2). In *Q. robur* seedlings, the effect of shortwave UV radiation (<350 nm) on adaxial epidermal flavonols content was even greater than the effect of blue light (Table 2).

Adaxial epidermal flavonol content was generally lower in the evergreen stands than in the deciduous stands, and the absolute (as opposed to relative) effects of our filter treatments were also smaller in the evergreen stands (Tables S16A-P, Fig. 4). Specifically, adaxial epidermal flavonol content was significantly lower in the evergreen stands for all those species found beneath both stand types (i.e. *A. platanoides, F. vesca*, and *O. acetosella*, Tables S16A-P, Fig. 4).

### Seasonal dynamics of anthocyanins are less responsive than flavonols to changes in light quality

The increase in adaxial epidermal anthocyanin content during understory leaf senescence was similar to the seasonal trend in flavonol content that we report. Although, unlike flavonols, adaxial epidermal anthocyanin content was not generally high during early spring. In particular, the largest increases in anthocyanin content during leaf senescence were present in plants whose leaves senesced latest in the year; *A. platanoides, A. podagraria* and *Q. robur* (Weeks 35-40, Fig. 5). Overall, seasonal changes in anthocyanin content in the adaxial epidermis of leaves were less responsive to our filter treatments than flavonol contents (Table 2, Figs. 4 & 5). Attenuating blue light significantly reduced anthocyanins in *A. platanoides* (Table 2, Tables S17A-D). This reduction in anthocyanin content in *A. platanoides* due to attenuating blue light, was greatest during the final weeks of autumn (17.7% reduction compared to the control, during week 38 averaged across stand types, including a 45.5% reduction in the evergreen stand during week 38, Fig. 5). However, attenuating blue light caused the opposite effect, increasing adaxial epidermal anthocyanin content by 7.57% in *M. bifolium* and 1.19% in *O. acetosella* (Tables S17I,J,M,N), though these effects were small (Table 2).

The effects of attenuating UV radiation below and above 350 nm were smaller than those of attenuating blue light on adaxial epidermal anthocyanin content (Table 2, Tables S17A-N, Figs. 4& 5). Attenuating UV radiation below 350 nm reduced anthocyanin content in *A. platanoides* and *A. podagraria* (Table 2, Tables S17A-D, Fig. 5), although only to a small degree (average across the measurement period by 8.9% and 9.8% respectively, Fig. 5). Attenuating UV radiation above 350nm had no significant effect on anthocyanin content (Tables S17A-N).

## Discussion

### Light quality has a small effect on spring phenology in the understory

Filter treatments revealed that the blue region of the solar spectrum can advance bud burst by an average of 1.4 days for *A. platanoides* seedlings growing in the forest understory, and final leaf out by 2.8 days (Fig. 2). Attenuating blue light from solar radiation in this sort of filter experiment (e.g. Siipola et al. 2015) also causes a reduction in PAR. However, the effect of blue light we report is consistent with findings from an experiment using branches of *B. pendula, Alnus glutinosa* and *Q. robur* under controlled conditions where PAR was equalised across filter treatments (Brelsford and Robson, 2018). Our study is also in line with a recent meta-analysis, in finding that UV radiation has a negligible effect on bud burst (Brelsford et al. 2019). Unlike *A. platanoides*, the emergence and leaf out of none of our herbaceous species responded to light quality in the understory. This confirms our hypothesis that leaf out in herbaceous species would be less responsive than tree seedlings to changes in light quality during spring. This result reaffirms the suggestion that the main cues controlling spring emergence in herbaceous species, which are submerged belowground during winter, are probably the timing of snow melt and increase in soil temperature rather than light quality (Price and Waser 1998, Rice et al. 2018). Herbaceous species were monitored less often and on a less detailed scale than *A. platanoides* meaning that subtle differences in their phenology could be overlooked. As such, increased sampling effort may be required to further test the effects of light quality on herbaceous species in future studies.

### Light quality affects autumn leaf senescence

Of those understory species with an autumn phenology which extends beyond canopy opening, the autumn leaf senescence of *A. platanoides, A. podagraria* and *R. cassubicus* was delayed by attenuating blue light. Blue light has been shown to enhance photosynthesis beyond simply its contribution to PAR, particularly in allowing the utilisation of transient sunflecks through rapid stomatal opening and rubisco activation (Sæbø et al. 1995, Goins et al. 1997; Matsuda et al. 2004; Košvancová-Zitová et al. 2009, Hogewoning et al. 2010). Attenuation of blue light throughout the growing season could reduce photosynthesis by impeding this rapid induction response, meaning that prolonged leaf retention is required to compensate for reduced carbon gain over the growing season (Chabot and Hicks, 1982, Zhang et al. 2013). As attenuating blue light from solar radiation also reduces PAR (by an average of 34.4 % in no blue no UV filter compared to control filter in our case), the delayed leaf senescence in response to attenuation of blue light that we report could be due to the reduction in PAR (Lee et al. 2003). This could also explain why attenuating blue light delayed leaf senescence the most in *A. platanoides* seedlings growing in the evergreen stand, where PAR was also lowest (Table S4, Fig. 3). However, neither our measurements of photosynthetic capacity nor stomatal conductance significantly differed across our filter treatments in *A. platanoides*, as might otherwise be expected in response to PAR or blue light (Fig. S8, Fig. S9). There is some evidence from research into other plant species, such as *Glycine max*, that the blue-light-detecting photoreceptors, cryptochromes, fulfill a specific role involving blue light in the acceleration of leaf senescence (Meng et al. 2013).

In contrast to *A. platanoides*, there was no effect of our filter treatments on the leaf senescence of *Q. robur* seedlings. Autumn leaf senescence in *Q. robur* occurred later and much more abruptly than that of the other species monitored. It appeared to coincide with a sudden drop in temperatures towards the end of autumn (week 40, Fig. 3, Fig S1). *Q. robur* may reabsorb less nutrients from its leaves compared to tree seedlings of similar species in the autumn (Hagen-Thorn et al. 2006). This suggestion is consistent with the peak of leaf chlorophyll content found during autumn canopy opening in *Q. robur* (Fig. S9). Furthermore, the photosynthetic capacity of *Q. robur* trees is slow to develop throughout the season (Morecroft et al. 2003) and could indicate that *Q. robur* has a different strategy involving extended carbon assimilation further into autumn compared to other species in our study.

As well as the effect of blue light, attenuating UV radiation below 350nm also delayed the onset of leaf senescence in *A. platanoides* (4.1 days delay for UV radiation below 350nm, and 14.3 days delay for blue light). This agrees with our hypothesis that attenuating UV radiation would delay leaf senescence. This is the first study to find that attenuating ambient solar UV radiation can delay autumn leaf senescence in understory tree seedlings. However, past research has found that supplemental UV-B radiation can accelerate leaf senescence in saplings of *Fagus sylvatica* growing in open-topped chambers compared with those in under simulated clear-sky solar UV-B radiation (Zeuthen et al. 1997). The delay in leaf senescence of *A. platanoides* under attenuated UV radiation below 350 nm could be attributable to leaves suffering less UV-induced photodamage (Zeuthen et al. 1997) throughout the growing season under this filter. However, if this was the mechanism responsible, it is unclear why only leaf senescence in *A. platanoides*, of the species we studied, responded in this way.

### Flavonol response was less sensitive to changes in light quality in shade-tolerant species than in more light-demanding species

We found that more light-demanding species tended to have the highest flavonol content throughout the season, and were more responsive to our filter treatments than most of the species with overwintering leaves (*O. acetosella* and *M. bifolium*) (Fig. 4). This confirms our hypothesis that shade-tolerant species would be less responsive to the filter treatments. In a study of tropical alpine plant types, no significant difference was found between the UV-screening in herbaceous species and woody species (Barnes et al. 2017). In contrast to this, a few comparative studies of plant types have suggested that the accumulation of UV-B absorbing compounds, is slightly more responsive to solar radiation in herbaceous species than in woody species, and that evergreen species are least responsive (Semerdjieva et al. 2003, Brzezinska and Kozlowska 2008, Li et al. 2010). These differences in flavonoid induction may be offset by higher constitutive contents of UV-B absorbing compounds in evergreen and woody species compared to herbaceous species (Semerdjieva et al. 2003, Li et al. 2010), possibly related to their greater leaf longevity, making investment in flavonoids worthwhile (Semerdjieva et al. 2003). However, we found that despite having shorter leaf longevity, the more light-demanding plants in our study had higher flavonol content throughout the season. Higher investment in flavonols by more light-demanding species may supplement other photoprotection mechanisms allowing species transient periods of high irradiance and canopy gaps to be exploited without severe photoinhibition (Takahashi and Badger, 2011). However, there were underlying seasonal trends in flavonol content under all our filter treatments, which implies that a variety of other factors, such as temperature and developmental stages of the plants, are also important determinants of leaf flavonol content (Pescheck and Bilger 2019, Hartikainen et al. 2019 submitted.

As we hypothesized, out of all our filter treatments, attenuation of blue light caused the largest reduction in adaxial epidermal flavonols in more light-demanding herbaceous species and deciduous tree seedlings (Fig. 4). These results are consistent with Siipola et al. (2015) who previously reported a larger effect of blue light than UV radiation on flavonol content in the pea, *Pisum sativum cv. Meteor* grown in full sunlight. Likewise, in a growth room, simulated understory blue light increased leaf adaxial epidermal flavonol content via cryptochrome photoreceptors under controlled PAR conditions (Brelsford et al. 2019). Accordingly, much of the reduction in flavonols during canopy closure during summer (Fig. 4), could be attributable to the reduction in blue light reaching the understory. Epidermal flavonols responded differently to our treatments in *Q. robur* compared with most other species. Attenuating UV radiation below 350 nm had the largest effect on *Q. robur* flavonols. This might suggest that at the core of is distribution range at lower latitudes where UV-B exposure is typically higher, UV-perception is more important as a cue (Bais et al. 2019).

The seasonal fluctuations in epidermal flavonols in the understory of the evergreen stands in our study, and the same stands in 2015-2016 (Hartikainen et al. 2019 submitted), were much smaller than those in the deciduous stands. Here, we found that the effects of attenuating blue light and UV radiation were also much smaller in the evergreen stands compared to the deciduous. While the lower flavonol contents in plants growing in the evergreen understory can largely be attributed to lower solar irradiance, and its blue and UV regions, the temperature differences between the stands are also likely to contribute to this effect (Davis et al. 2019, Pescheck and Bilger 2019). Evergreen stands are warmer during the winter and cooler during the summer, because of the insulating effects of a dense closed canopy all year round (Davis et al. 2019). We should also note that ventilation gaps meant that our filter treatments did not totally exclude the attenuated regions of radiation, and the small percentages of solar blue and UV radiation reaching the plants could have had an effect.

### Light quality affects flavonol and anthocyanin content in autumn

Until recently, leaf flavonol contents were generally considered not to increase during autumn, because of the presumed inefficiency of investing in leaves at the end of their life (reviewed by Wilkinson et al. 2002). However, Mattila et al. (2018) found that flavonols increase prior to and during autumn senescence in the tree species *B. pendula* and *Sorbus aucuparia*. Accordingly, we report an autumn increase in adaxial epidermal flavonol content in *A. platanoides* and *Q. robur* seedlings, as well as understory *A. podagraria*, and wintergreen *F. vesca* (Weeks 35-40). This increase in flavonols following canopy opening was much smaller, or absent, in plants under filters attenuating UV radiation and blue light (Weeks 35-40, Fig. 3). In autumn, adaxial epidermal anthocyanin content like that of epidermal flavonols tended to increase. However, anthocyanins generally responded less strongly than flavonols to attenuation of blue light and UV radiation.

It is possible that the increase in flavonols and anthocyanins during autumn, is due to the increase in solar radiation reaching the understory after canopy opening (Richardson and O’Keefe 2009), including shortwave radiation in the blue and UV regions (Hoch et al. 2001). However, there was also an autumnal increase in flavonol and anthocyanin content for species growing in the evergreen stand, in the absence of canopy opening (Ross and Flanagan, 1986, Constabel and Lieffers, 1996). This suggests that other factors such as colder temperatures during autumn also contribute to the autumn increase in flavonol and anthocyanin content (Pescheck and Bilger 2019, Renner and Zohner, 2019). Equally, our results also validate the hypothesis that the increase in anthocyanins may be a part of the leaf senescence timetable, and supports their role as antioxidants to remove ROS produced during chlorophyll breakdown (Archetti et al. 2009).

## Conclusions

Both blue light and UV radiation affected the phenology and leaf pigmentation of understory plants. In general, flavonols were most responsive to change in blue light, which makes up a larger portion of the spectrum than UV radiation even in canopy shade. Bud burst was only advanced by blue light in *A. platanoides* seedlings, whose leaf senescence was hastened by UV radiation in the understory. Whereas, blue light delayed autumn leaf senescence in most of the understory species. An extended period of canopy shading into autumn, and thus reduced blue light and UV radiation in the understory, is predicted under climate warming (Piao et al. 2019, Vitasse et al. 2011, Buitenwerf et al. 2015). Considered together with our results, this implies that the growing season in the understory would be extended for most plants that senesce in autumn. As such, future research should investigate how changes in blue and UV radiation will produce phenological mismatches in the understory plant community because of their differential effects on different plant functional types.

## Acknowledgements

This research was funded by decisions #266523 and #304519 to TMR from the Academy of Finland, the Doctoral Programme in Plant Science grant and Lammi Biological Station’s Environmental Research Foundation grant 2017 and 2018 to CCB. We thank Lammi Biological Station, Paula Lebowski, Santa Neimane, and Marta Pieristè for the help with fieldwork, and Titta Kotilainen for her advice on plotting spectra.

