## Supplementary materials for "Understory light quality affects leaf pigments and leaf phenology in different plant functional types"

Table S1. Spectral irradiance for different regions and ratios of the solar spectrum, in different forest stands across different dates.

| Date | stand | TYPE | PAR(PPFD) | UV-B(q) | UV-A(q) | Blue(q) | Green(q) | Red(q) | Far-red(q) | UV-B:PAR | R:FR | B:G |
| --- | --- | --- | --- | --- | --- | --- | --- | --- | --- | --- | --- | --- |
| 2015/04/25 | Betula Old | sun | 882.01 | 0.60 | 61.33 | 171.36 | 214.48 | 202.95 | 169.82 | 0.72 | 1.03 | 0.80 |
| 2015/04/25 | Betula Young | sun | 963.39 | 0.78 | 68.98 | 187.93 | 234.01 | 220.80 | 184.84 | 0.83 | 1.03 | 0.80 |
| 2015/04/25 | Betula Old | sun | 86.42 | 0.06 | 6.53 | 17.11 | 21.20 | 19.42 | 19.08 | 0.84 | 0.82 | 0.81 |
| 2015/04/25 | Quercus | sun | 1095.77 | 0.72 | 74.85 | 212.52 | 266.88 | 251.82 | 209.18 | 0.67 | 1.04 | 0.80 |
| 2015/05/22 | Betula Old | sun | 729.89 | 0.58 | 50.04 | 139.58 | 180.12 | 166.65 | 150.14 | 0.88 | 0.92 | 0.78 |
| 2015/05/22 | Betula Young | sun | 1043.08 | 0.88 | 68.66 | 197.39 | 258.05 | 238.81 | 218.31 | 0.85 | 0.93 | 0.76 |
| 2015/05/22 | P. abies | sun | 521.44 | 0.34 | 32.76 | 99.37 | 127.40 | 120.81 | 102.11 | 0.66 | 1.01 | 0.78 |
| 2015/05/22 | Quercus | sun | 1049.05 | 0.99 | 76.37 | 205.79 | 256.66 | 237.81 | 197.89 | 0.95 | 1.03 | 0.80 |
| 2015/07/05 | Betula Old | sun | 570.20 | 0.44 | 35.51 | 107.38 | 141.09 | 131.27 | 123.61 | 0.76 | 0.84 | 0.76 |
| 2015/07/05 | Betula Young | sun | 746.76 | 0.47 | 46.37 | 140.97 | 184.46 | 172.30 | 162.10 | 0.63 | 0.89 | 0.76 |
| 2015/07/05 | P. abies | sun | 871.34 | 0.53 | 52.84 | 165.20 | 213.09 | 202.49 | 166.01 | 0.64 | 1.05 | 0.78 |
| 2015/07/05 | Quercus | sun | 756.19 | 0.46 | 46.81 | 141.70 | 187.45 | 174.26 | 163.35 | 0.61 | 0.89 | 0.76 |
| 2015/08/21 | Betula Old | sun | 286.61 | 0.23 | 19.09 | 54.92 | 71.05 | 65.26 | 65.67 | 0.87 | 0.82 | 0.77 |
| 2015/08/21 | Betula Young | sun | 814.06 | 0.43 | 49.53 | 153.27 | 200.55 | 188.62 | 167.27 | 0.53 | 0.98 | 0.76 |
| 2015/08/21 | Quercus | sun | 561.53 | 0.47 | 35.39 | 105.99 | 138.33 | 129.84 | 118.32 | 0.83 | 0.95 | 0.77 |
| 2015/04/25 | Betula Old | shade | 144.05 | 0.34 | 22.47 | 35.02 | 34.41 | 27.70 | 26.72 | 2.35 | 0.87 | 1.02 |
| 2015/04/25 | Betula Young | shade | 206.41 | 0.42 | 26.10 | 45.90 | 49.49 | 42.93 | 39.96 | 2.11 | 0.90 | 0.94 |
| 2015/04/25 | P. abies | shade | 20.36 | 0.04 | 2.86 | 4.65 | 5.02 | 4.00 | 6.33 | 2.01 | 0.50 | 0.93 |
| 2015/04/25 | Quercus | shade | 190.61 | 0.32 | 24.79 | 43.78 | 46.02 | 38.47 | 33.77 | 1.68 | 0.98 | 0.96 |
| 2015/05/22 | Betula Old | shade | 85.15 | 0.26 | 14.03 | 19.29 | 22.71 | 14.74 | 29.07 | 3.05 | 0.38 | 0.85 |
| 2015/05/22 | Betula Young | shade | 111.83 | 0.24 | 13.55 | 21.78 | 30.60 | 21.35 | 44.87 | 2.10 | 0.35 | 0.71 |
| 2015/05/22 | P. abies | shade | 29.90 | 0.04 | 3.43 | 6.44 | 7.41 | 6.15 | 9.52 | 1.49 | 0.50 | 0.88 |
| 2015/05/22 | Quercus | shade | 93.13 | 0.42 | 19.69 | 24.66 | 23.37 | 14.90 | 22.01 | 4.53 | 0.52 | 1.05 |
| 2015/07/05 | Betula Old | shade | 49.07 | 0.17 | 7.79 | 10.74 | 13.87 | 8.24 | 28.23 | 3.43 | 0.20 | 0.77 |
| 2015/07/05 | Betula Young | shade | 40.21 | 0.10 | 6.00 | 8.19 | 11.85 | 6.73 | 29.69 | 2.66 | 0.15 | 0.69 |
| 2015/07/05 | P. abies | shade | 17.61 | 0.03 | 2.68 | 4.06 | 4.44 | 3.32 | 6.46 | 3.15 | 0.32 | 0.98 |
| 2015/07/05 | Quercus | shade | 51.45 | 0.13 | 7.32 | 9.41 | 15.47 | 8.64 | 31.05 | 2.44 | 0.19 | 0.61 |
| 2015/08/21 | Betula Old | shade | 17.30 | 0.06 | 3.10 | 3.98 | 5.11 | 2.59 | 15.54 | 3.61 | 0.12 | 0.77 |
| 2015/08/21 | Betula Young | shade | 38.74 | 0.08 | 5.01 | 8.34 | 10.52 | 7.11 | 24.29 | 1.97 | 0.21 | 0.79 |
| 2015/08/21 | Quercus | shade | 19.69 | 0.07 | 3.30 | 4.25 | 6.01 | 2.95 | 18.86 | 3.78 | 0.11 | 0.71 |
| 2015/05/22 | Betula Old | leaf | 391.82 | 0.43 | 31.41 | 76.52 | 97.53 | 87.01 | 86.05 | 1.12 | 0.84 | 0.79 |
| 2015/05/22 | Betula Young | leaf | 557.49 | 0.55 | 40.17 | 105.89 | 139.36 | 125.46 | 127.66 | 1.01 | 0.80 | 0.76 |
| 2015/05/22 | P. abies | leaf | 324.60 | 0.16 | 20.84 | 62.50 | 79.49 | 74.52 | 64.30 | 0.52 | 0.98 | 0.79 |
| 2015/05/22 | Betula Old | leaf | 627.18 | 0.59 | 43.36 | 120.92 | 155.45 | 141.89 | 135.78 | 1.03 | 0.81 | 0.78 |
| 2015/07/05 | Betula Young | leaf | 199.24 | 0.20 | 15.07 | 38.13 | 50.71 | 43.94 | 59.11 | 1.21 | 0.54 | 0.75 |
| 2015/07/05 | P. abies | leaf | 112.52 | 0.07 | 8.25 | 21.95 | 27.61 | 25.48 | 24.30 | 1.29 | 0.72 | 0.83 |
| 2015/07/05 | Quercus | leaf | 304.50 | 0.26 | 21.78 | 56.90 | 77.11 | 68.18 | 78.30 | 0.95 | 0.65 | 0.72 |
| 2015/08/21 | Betula Old | leaf | 149.75 | 0.19 | 11.61 | 29.17 | 37.39 | 33.42 | 40.38 | 1.51 | 0.64 | 0.79 |
| 2015/08/21 | Betula Young | leaf | 140.86 | 0.03 | 9.33 | 26.67 | 35.79 | 31.43 | 43.50 | 0.24 | 0.59 | 0.75 |
| 2015/08/21 | Quercus | leaf | 207.29 | 0.19 | 14.51 | 39.61 | 51.94 | 46.68 | 53.32 | 0.98 | 0.65 | 0.75 |
| 2016/04/21 | Picea | leaf | 29.01 | 0.04 | 3.08 | 6.18 | 7.12 | 6.06 | 7.73 | 2.30 | 0.50 | 0.95 |
| 2016/04/22 | Picea | leaf | 820.64 | 0.59 | 56.77 | 158.26 | 198.64 | 188.35 | 160.14 | 0.76 | 0.95 | 0.80 |
| 2016/05/09 | Betula Old | leaf | 292.21 | 0.44 | 26.07 | 59.05 | 72.52 | 63.14 | 60.70 | 1.53 | 0.85 | 0.82 |
| 2016/05/09 | Betula Young | leaf | 263.14 | 0.42 | 22.20 | 50.59 | 66.92 | 56.92 | 66.33 | 1.69 | 0.65 | 0.76 |
| 2016/05/09 | Picea | leaf | 17.45 | 0.07 | 3.00 | 4.57 | 4.42 | 2.86 | 5.13 | 3.91 | 0.40 | 1.04 |
| 2016/05/09 | Quercus | leaf | 395.78 | 0.57 | 33.64 | 79.00 | 96.72 | 87.73 | 77.23 | 1.61 | 0.92 | 0.83 |
| 2016/05/24 | BetulaOld | leaf | 79.49 | 0.21 | 9.43 | 16.71 | 21.04 | 15.37 | 28.44 | 2.89 | 0.38 | 0.80 |
| 2016/05/24 | Picea | leaf | 25.38 | 0.07 | 3.43 | 5.74 | 6.18 | 5.06 | 6.07 | 4.49 | 0.59 | 0.90 |
| 2016/05/25 | BetulaYoung | leaf | 70.96 | 0.12 | 6.63 | 13.93 | 19.05 | 14.32 | 30.22 | 1.97 | 0.31 | 0.72 |
| 2016/05/25 | Quercus | leaf | 62.81 | 0.14 | 6.43 | 11.90 | 18.26 | 11.52 | 33.37 | 2.22 | 0.23 | 0.65 |
| 2016/04/21 | BetulaOld | shade | 99.63 | 0.28 | 18.29 | 26.30 | 23.73 | 17.29 | 18.15 | 2.76 | 0.74 | 1.11 |
| 2016/04/21 | BetulaYoung | shade | 114.31 | 0.35 | 20.72 | 29.31 | 27.09 | 20.54 | 21.60 | 3.10 | 0.74 | 1.08 |
| 2016/04/21 | Picea | shade | 8.86 | 0.03 | 1.96 | 2.40 | 2.21 | 1.37 | 3.83 | 3.80 | 0.25 | 1.09 |
| 2016/04/21 | Quercus | shade | 94.56 | 0.33 | 19.44 | 26.08 | 22.40 | 15.56 | 16.53 | 3.53 | 0.72 | 1.16 |
| 2016/05/09 | Betula Old | shade | 138.91 | 0.37 | 18.85 | 32.72 | 35.60 | 25.22 | 30.22 | 2.64 | 0.65 | 0.92 |
| 2016/05/09 | Betula Young | shade | 112.76 | 0.35 | 15.03 | 24.46 | 30.58 | 20.13 | 37.01 | 3.10 | 0.39 | 0.80 |
| 2016/05/09 | Picea | shade | 16.23 | 0.07 | 3.01 | 4.27 | 4.10 | 2.63 | 4.97 | 4.27 | 0.38 | 1.04 |
| 2016/05/09 | Quercus | shade | 132.60 | 0.45 | 21.85 | 34.19 | 33.14 | 22.53 | 24.04 | 3.39 | 0.73 | 1.03 |
| 2016/05/24 | BetulaOld | shade | 54.63 | 0.19 | 7.70 | 11.89 | 15.15 | 9.55 | 25.50 | 3.48 | 0.26 | 0.78 |
| 2016/05/24 | Picea | shade | 12.72 | 0.05 | 2.08 | 3.14 | 3.31 | 2.13 | 4.93 | 3.78 | 0.31 | 0.95 |
| 2016/05/25 | BetulaYoung | shade | 37.07 | 0.09 | 4.73 | 7.85 | 10.77 | 6.26 | 23.27 | 2.51 | 0.18 | 0.73 |
| 2016/05/25 | Quercus | shade | 49.93 | 0.13 | 5.82 | 9.73 | 14.99 | 8.47 | 29.96 | 2.61 | 0.19 | 0.65 |
| 2016/04/21 | BetulaOld | sun | 472.38 | 0.38 | 36.80 | 94.03 | 114.23 | 106.23 | 91.95 | 0.95 | 0.92 | 0.84 |
| 2016/04/21 | BetulaYoung | sun | 803.60 | 0.62 | 57.14 | 155.88 | 194.48 | 183.62 | 156.96 | 0.78 | 0.95 | 0.80 |
| 2016/04/21 | Picea | sun | 111.58 | 0.06 | 7.33 | 21.39 | 27.20 | 25.55 | 23.81 | 1.22 | 0.72 | 0.85 |
| 2016/04/21 | Quercus | sun | 849.79 | 0.61 | 58.58 | 164.07 | 205.82 | 194.83 | 165.26 | 0.74 | 0.95 | 0.80 |
| 2016/04/29 | Picea | sun | 602.18 | 0.31 | 31.94 | 109.44 | 146.48 | 142.95 | 119.43 | 0.51 | 0.99 | 0.75 |
| 2016/05/09 | Betula Old | sun | 664.57 | 0.56 | 41.62 | 121.28 | 162.32 | 156.14 | 136.64 | 0.85 | 0.96 | 0.75 |
| 2016/05/09 | Betula Young | sun | 848.53 | 0.76 | 51.85 | 154.41 | 209.06 | 197.96 | 179.73 | 0.91 | 0.90 | 0.74 |
| 2016/05/09 | Picea | sun | 485.77 | 0.30 | 25.68 | 86.50 | 117.83 | 116.88 | 96.65 | 0.62 | 1.01 | 0.73 |
| 2016/05/09 | Quercus | sun | 843.52 | 0.72 | 53.91 | 156.65 | 204.97 | 197.68 | 164.99 | 0.87 | 1.00 | 0.77 |
| 2016/05/24 | BetulaOld | sun | 672.95 | 0.57 | 41.08 | 125.14 | 165.78 | 155.75 | 138.96 | 0.86 | 0.92 | 0.75 |
| 2016/05/24 | Picea | sun | 485.89 | 0.38 | 28.56 | 90.24 | 118.83 | 113.39 | 92.28 | 0.77 | 1.04 | 0.76 |
| 2016/05/25 | BetulaYoung | sun | 510.21 | 0.31 | 27.23 | 90.84 | 125.68 | 121.35 | 113.55 | 0.65 | 0.86 | 0.72 |

Table S2. LAI of different plots and stands taken on 09/06/2017.

| **Stand** | **Plot** | **Date** | **LAI.se** | **LAI.mean** |
| --- | --- | --- | --- | --- |
| Betula pendula old | 1 | 09/06/2017 | 0.0066667 | 1.696667 |
| Betula pendula old | 2 | 09/06/2017 | 0.0233333 | 1.983333 |
| Betula pendula old | 3 | 09/06/2017 | 0.0517472 | 2.063333 |
| Betula pendula young | 1 | 09/06/2017 | 0.0732575 | 2.48 |
| Betula pendula young | 2 | 09/06/2017 | 0.1068228 | 2.343333 |
| Betula pendula young | 3 | 09/06/2017 | 0.0808977 | 2.273333 |
| Picea abies | 1 | 09/06/2017 | 0.1369104 | 3.696667 |
| Picea abies | 2 | 09/06/2017 | 0.1159023 | 3.66 |
| Picea abies | 3 | 09/06/2017 | 0.1223837 | 3.833333 |
| Picea abies | 4 | 09/06/2017 | 0.0945163 | 3.32 |
| Picea abies | 5 | 09/06/2017 | 0.117804 | 3.203333 |
| Picea abies | 6 | 09/06/2017 | 0.1244097 | 3.406667 |
| Quercus robur | 1 | 09/06/2017 | 0.0057735 | 1.5 |
| Quercus robur | 2 | 09/06/2017 | 0.0057735 | 1.53 |
| Quercus robur | 3 | 09/06/2017 | 0.0470225 | 1.453333 |

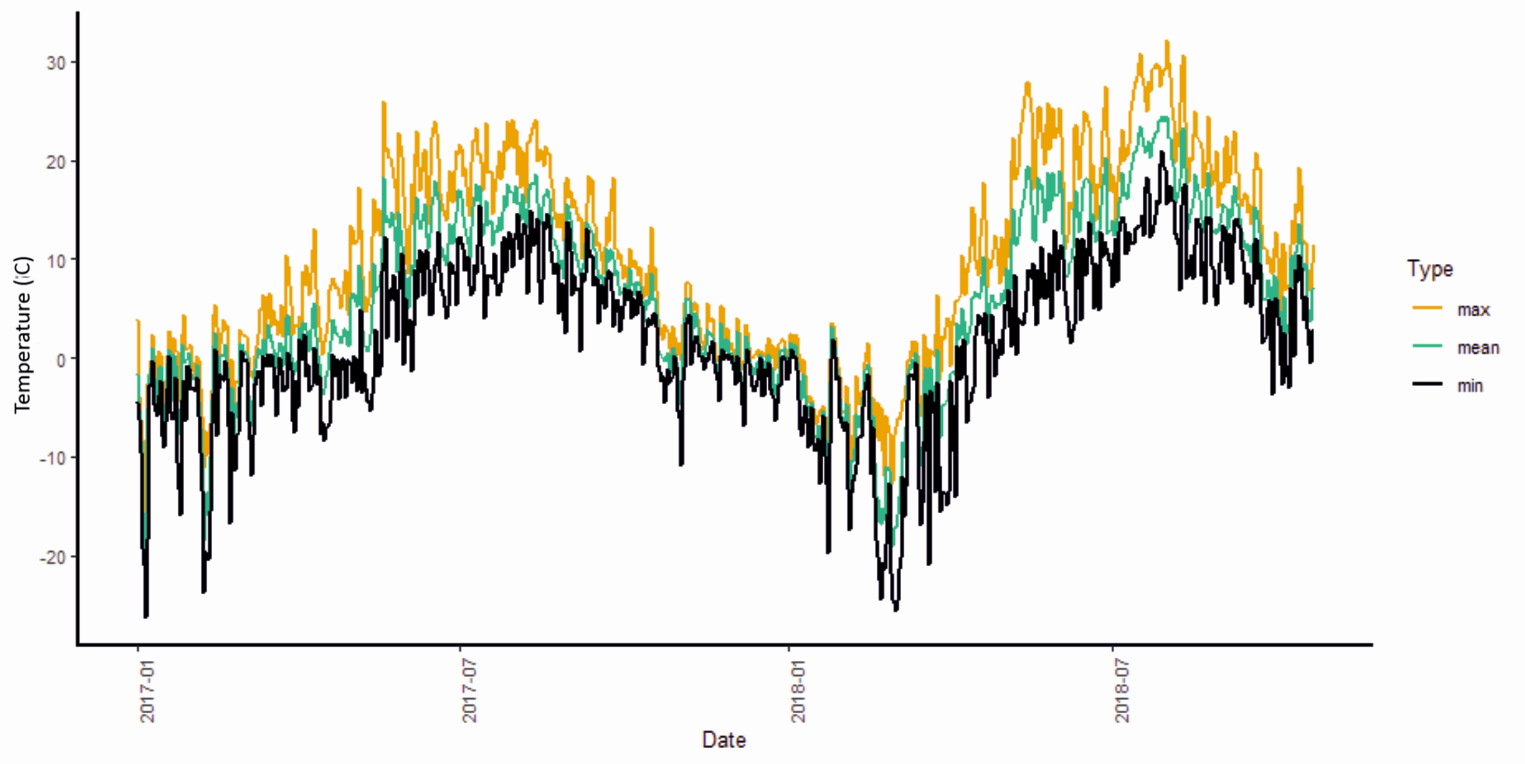

Figure S1. Ambient temperature (not in forest) 2017-2018 for Lammi Biological station. The data was recorded at the site and processed by the Finnish Meteorological Institute (<https://en.ilmatieteenlaitos.fi/weather/h%C3%A4meenlinna/lammi>).

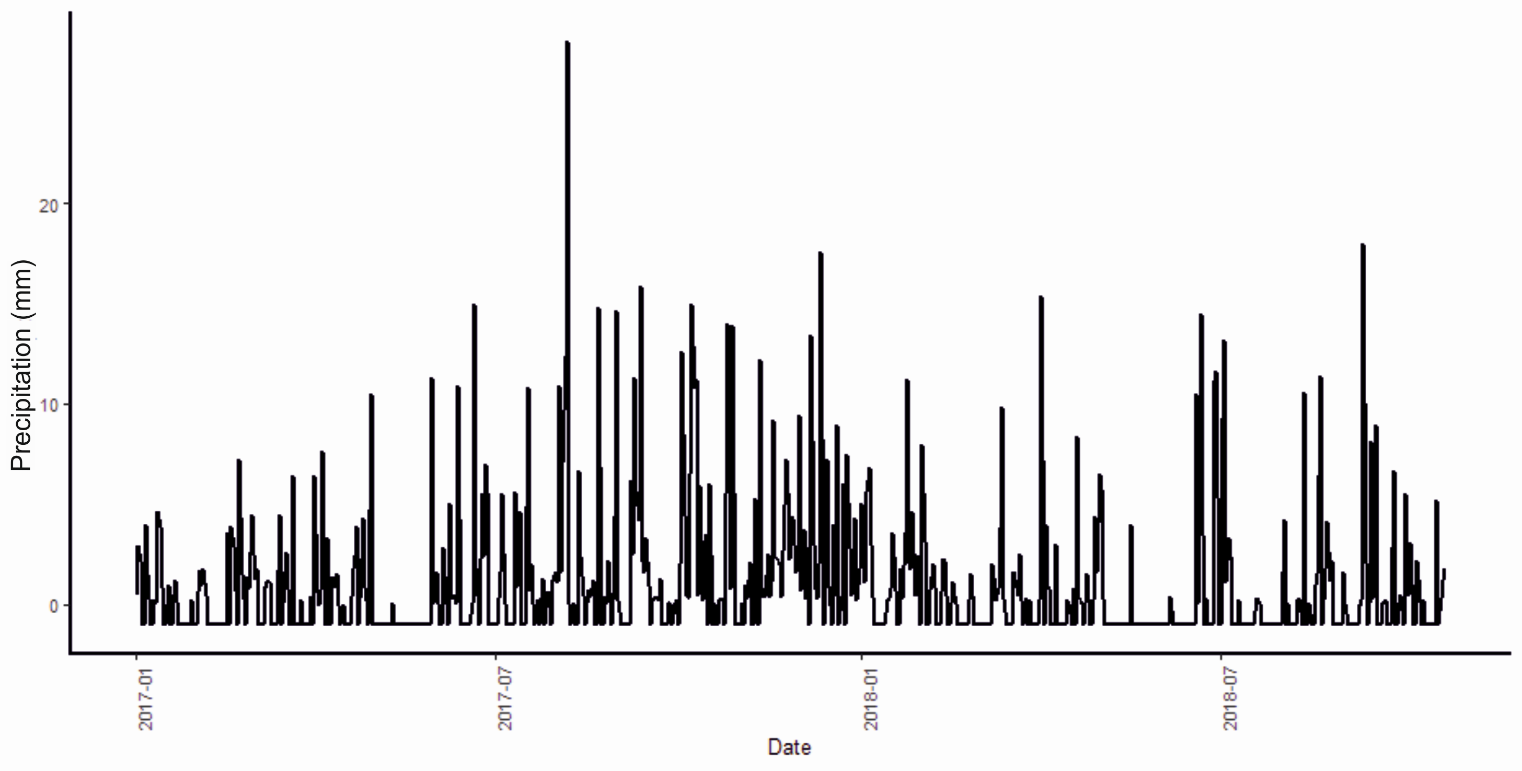

Figure S2. Precipitation 2017-2018 for Lammi Biological station. The data was recorded at the site and processed by the Finnish Meteorological Institute (<https://en.ilmatieteenlaitos.fi/weather/h%C3%A4meenlinna/lammi>).

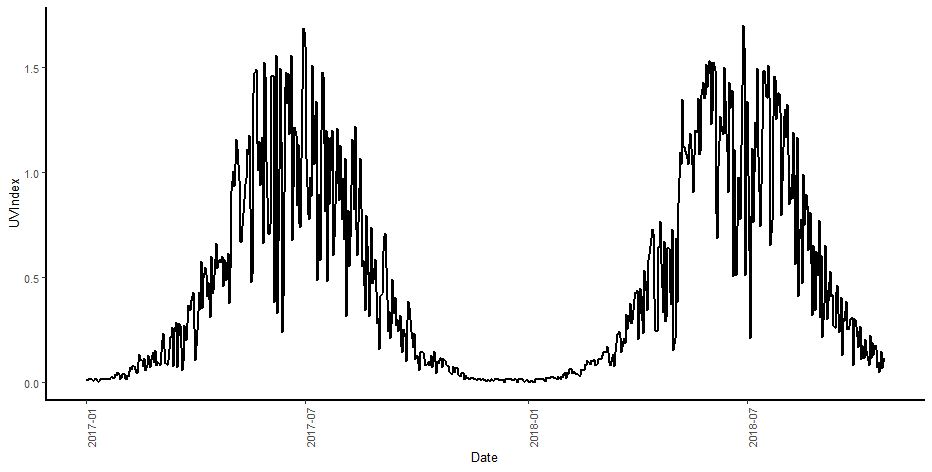

Figure S3. UV Index 2017-2018 from FMI. <https://en.ilmatieteenlaitos.fi/download-observations#!/> Taken from Jokioinen Ilmala, which is the closest station to the study site in Lammi with UV irradiance data.

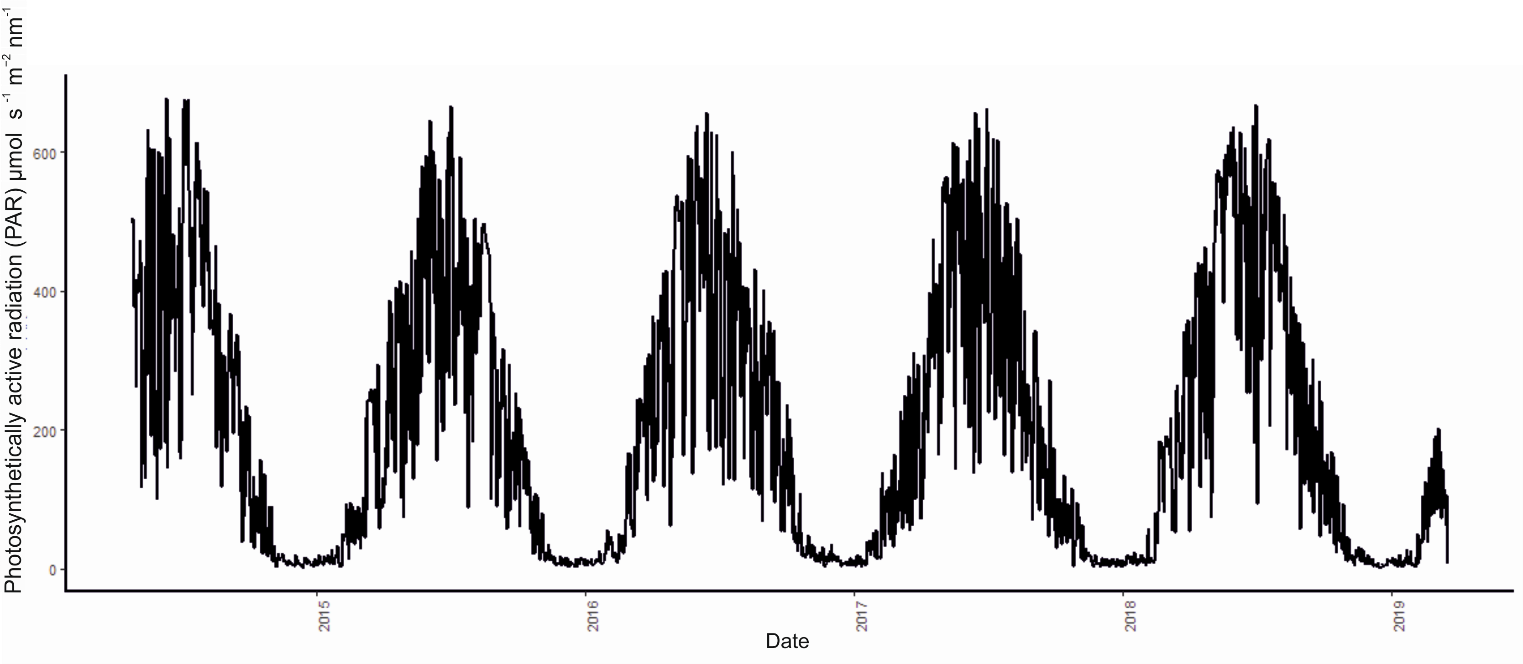

Figure S4. Ambient PAR (not in forest) 2015-2019 Ambient solar PAR above the canopy at Lammi Biological Station between 2015-2019 was also recorded to show the effects of changing cloudiness on incoming irradiance (PQS1 PAR Quantum Sensor, Kipp & Zonen, Delft, Netherlands, Fig. S4).

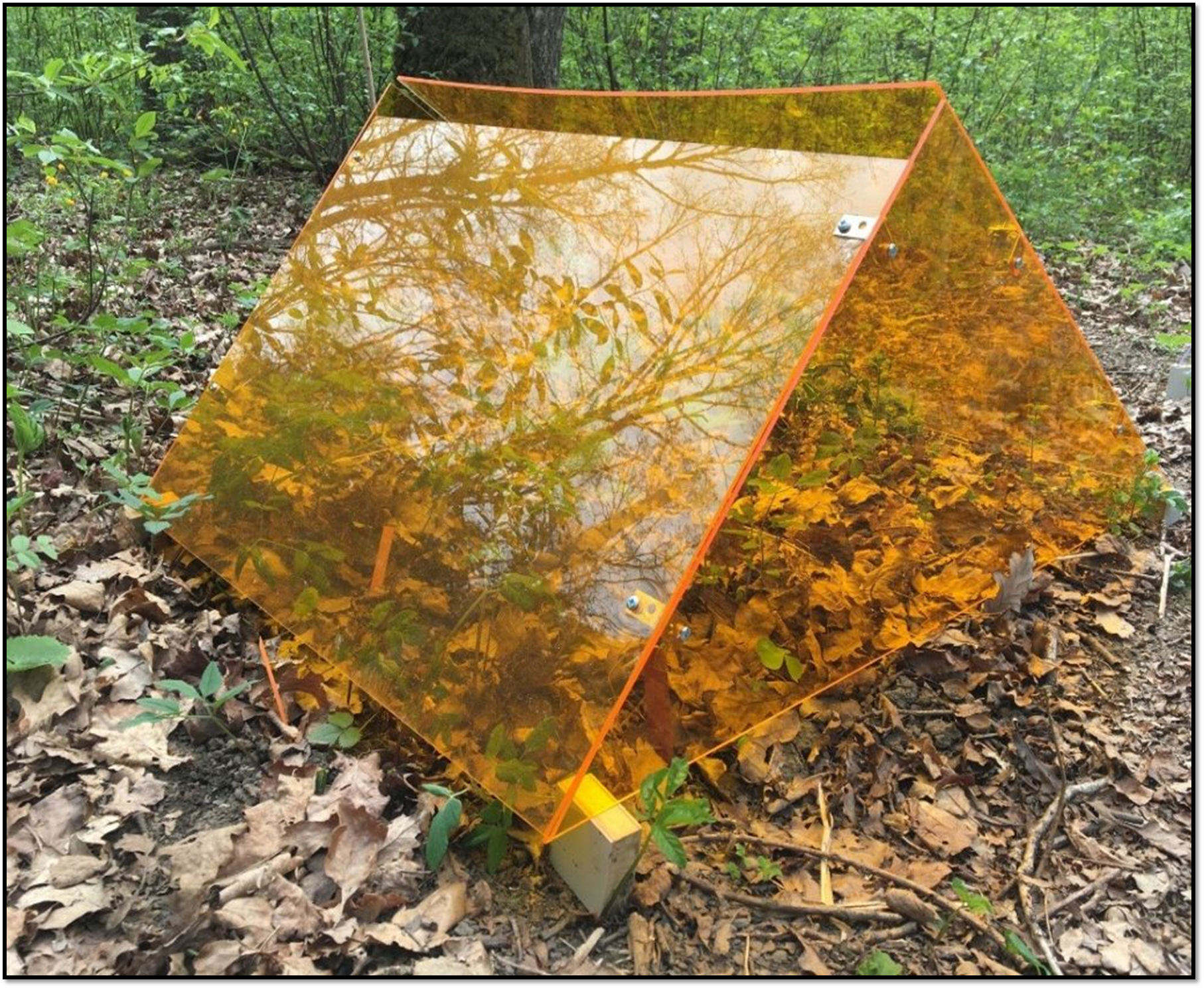

Fig. S5. Example of the filter structure in the field. This filter is a “No blue and UV treatment” installed in a deciduous stand with a *Q. robur* canopy. All filter treatments had identical filter structures (88 cm length x 60 cm width x 40 cm in height) with ventilation slots at the apex and around the base.

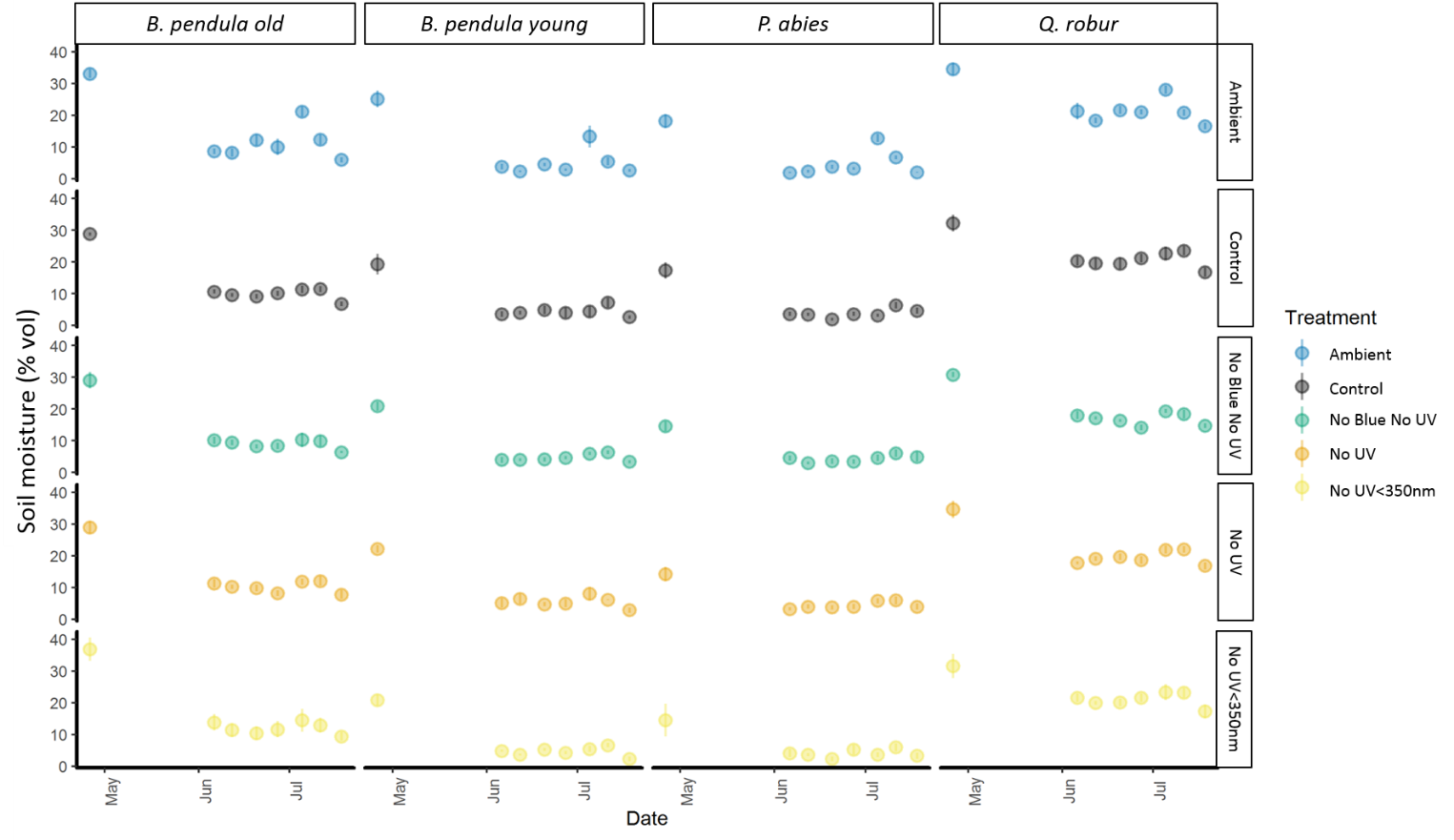

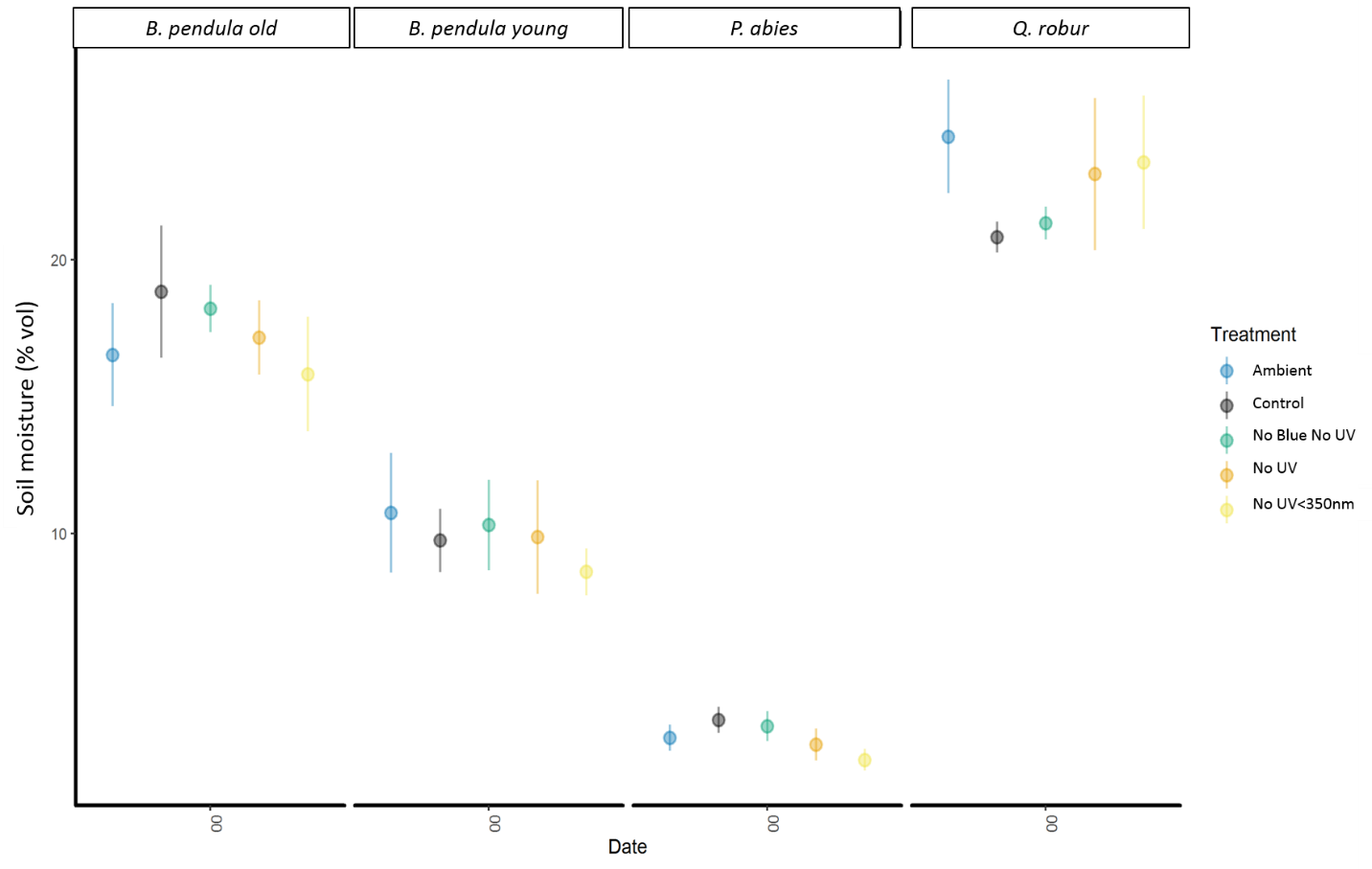

Figure S6. Soil moisture % vol shown for ambient soil conditions, and under different filter treatments in the forest understory of different stands. The above graph shows soil moisture during 2018, whilst the below measurement was taken on 17^th^ June 2017.

A
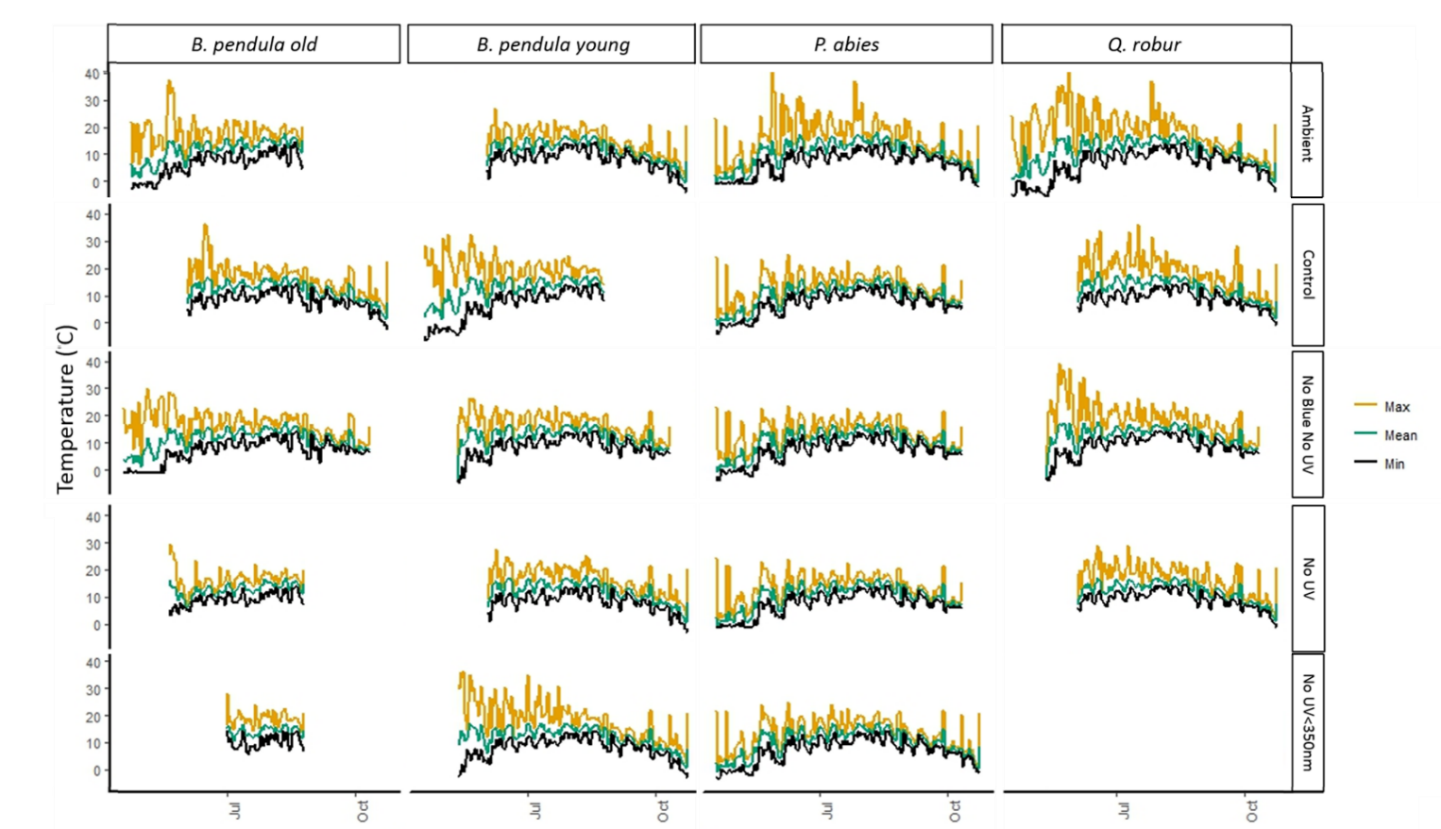
B

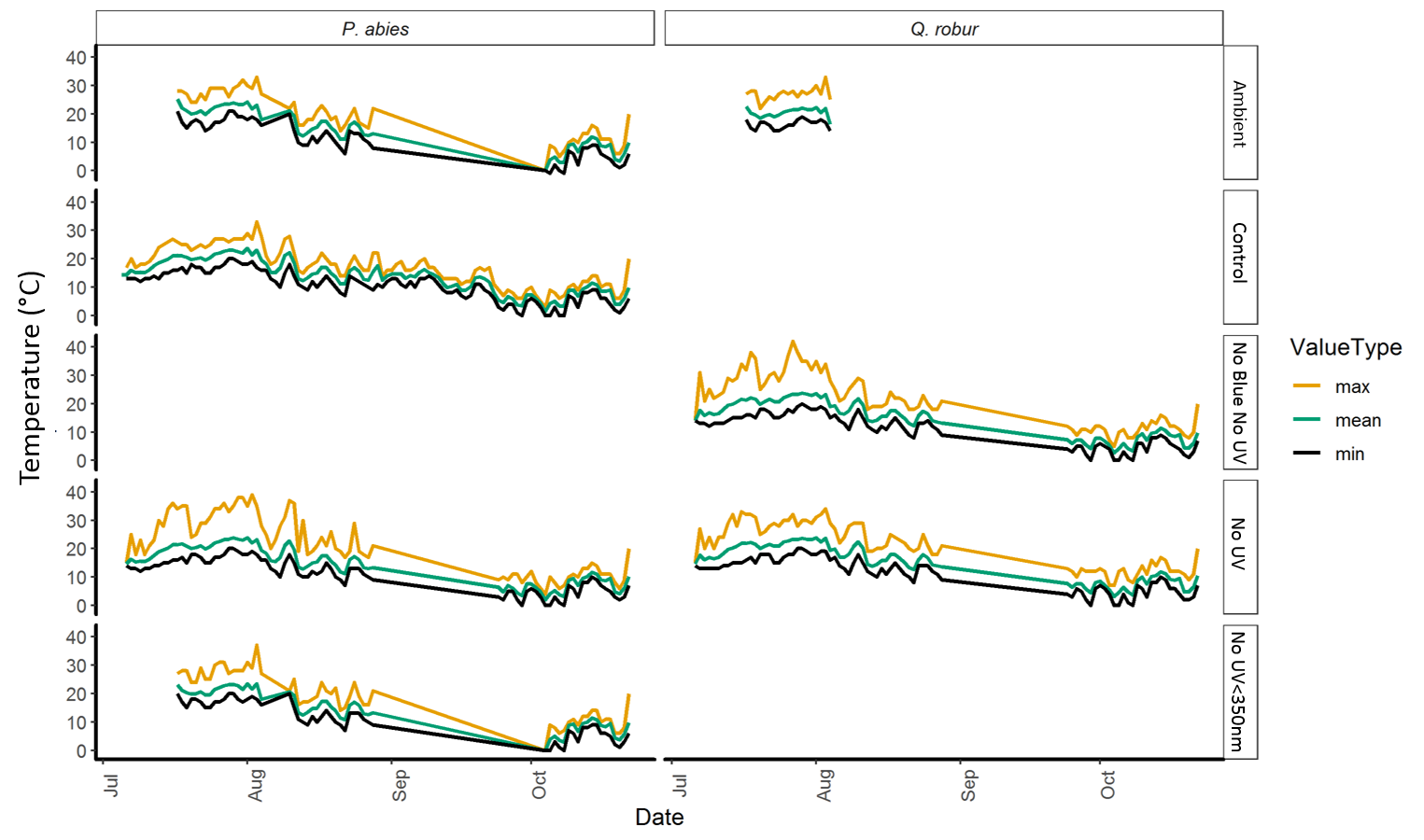
Figure S7 (A-B). Understory different filters and under different stands. A) Shown above is the temperature during 2017, and, B) below = 2018. Interpolated data for 2018, between September and October, shown in Figure S6C.

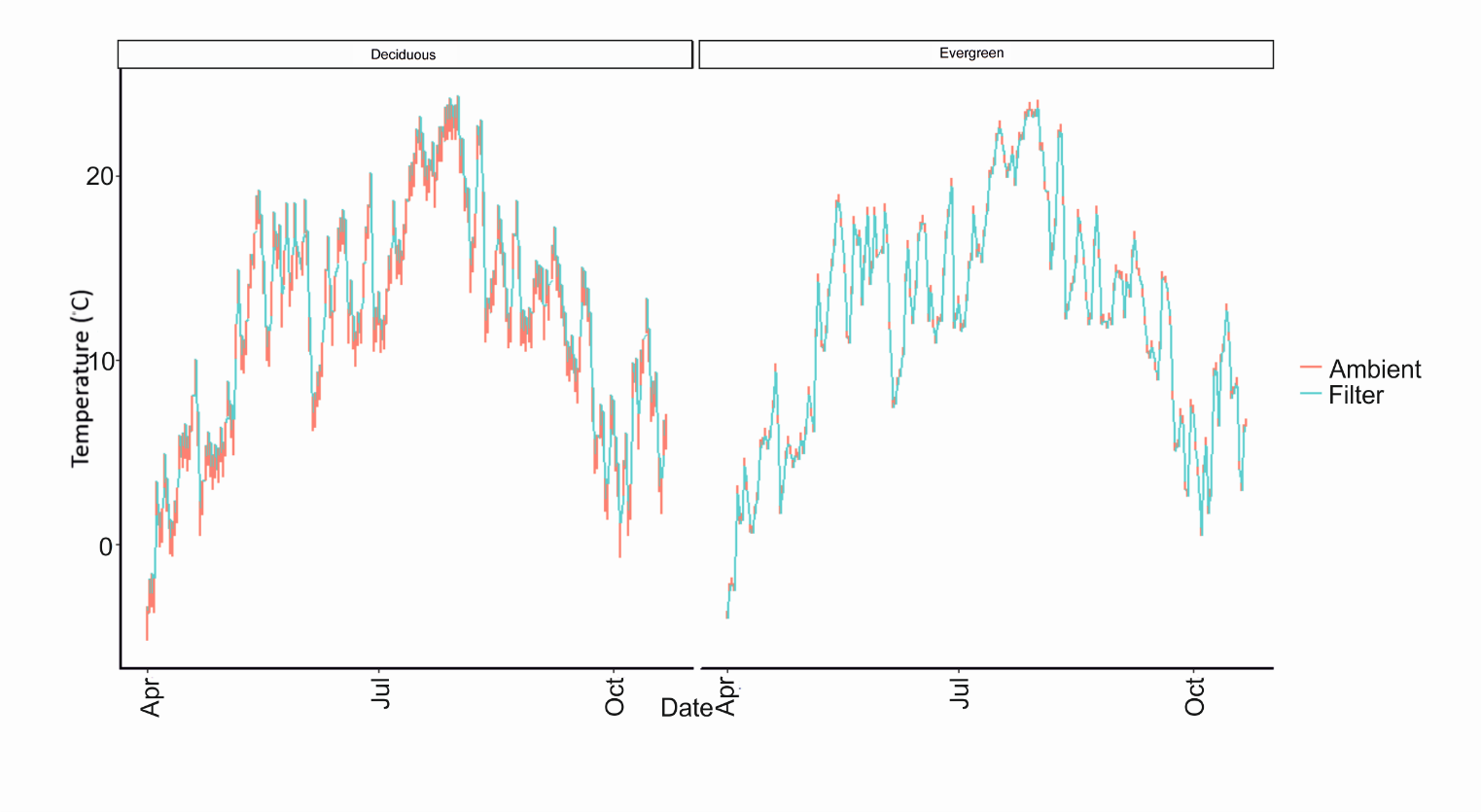

Figure S8C. Ambient interpolated temperature data for 2018 shown here. Data was calculated based on ambient temperature outside the forest canopy, and the filter temperature based on the mean average difference between filter temperature and outside temperature.

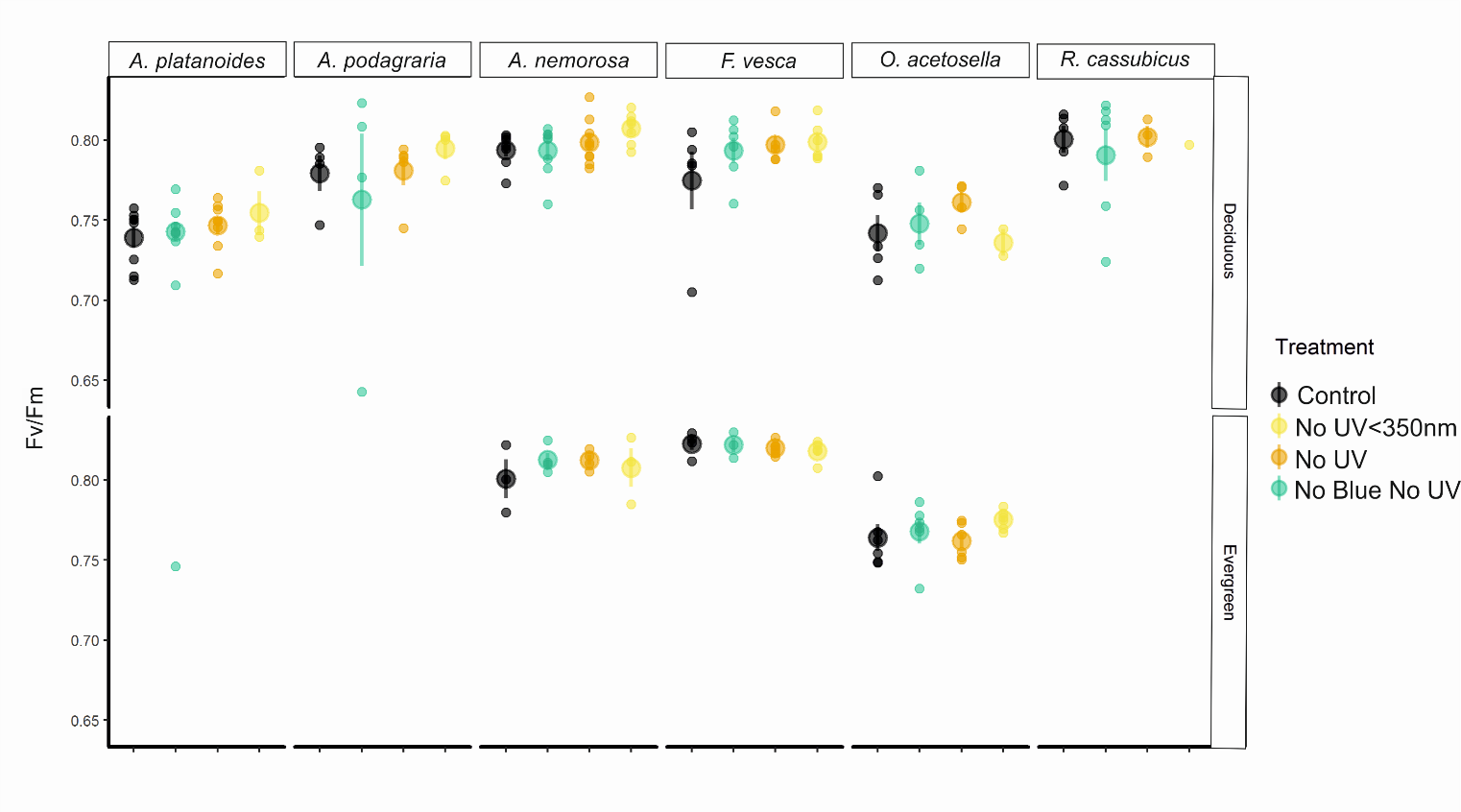

Figure S9. Fv/Fm of different understorey plants measured on 24^th^ -25th June 2017 between 10:00-16:00.

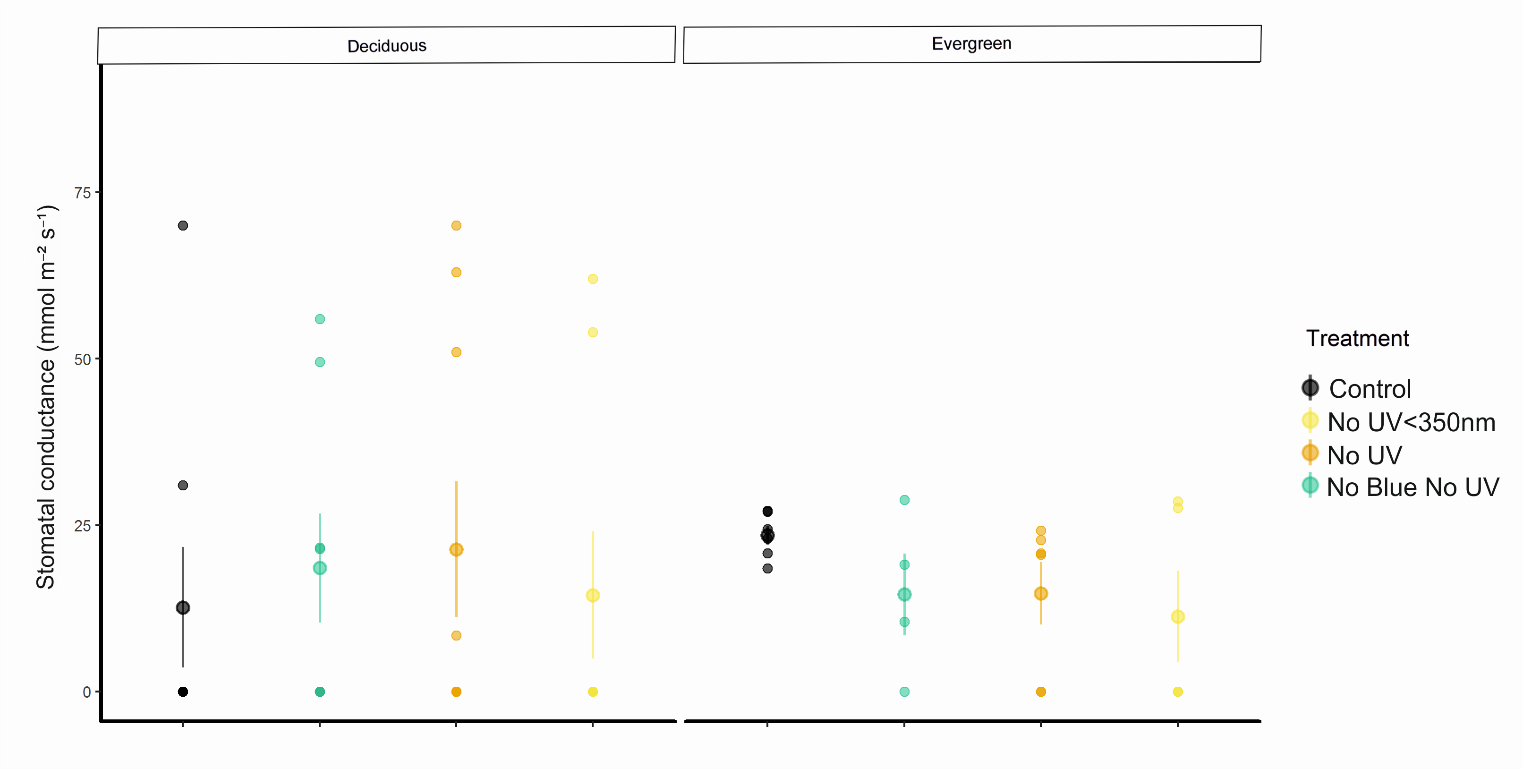

Figure S10. Stomatal conductance of *A. platanoides* on 16^th^ June between 10:00-14:00 2018.

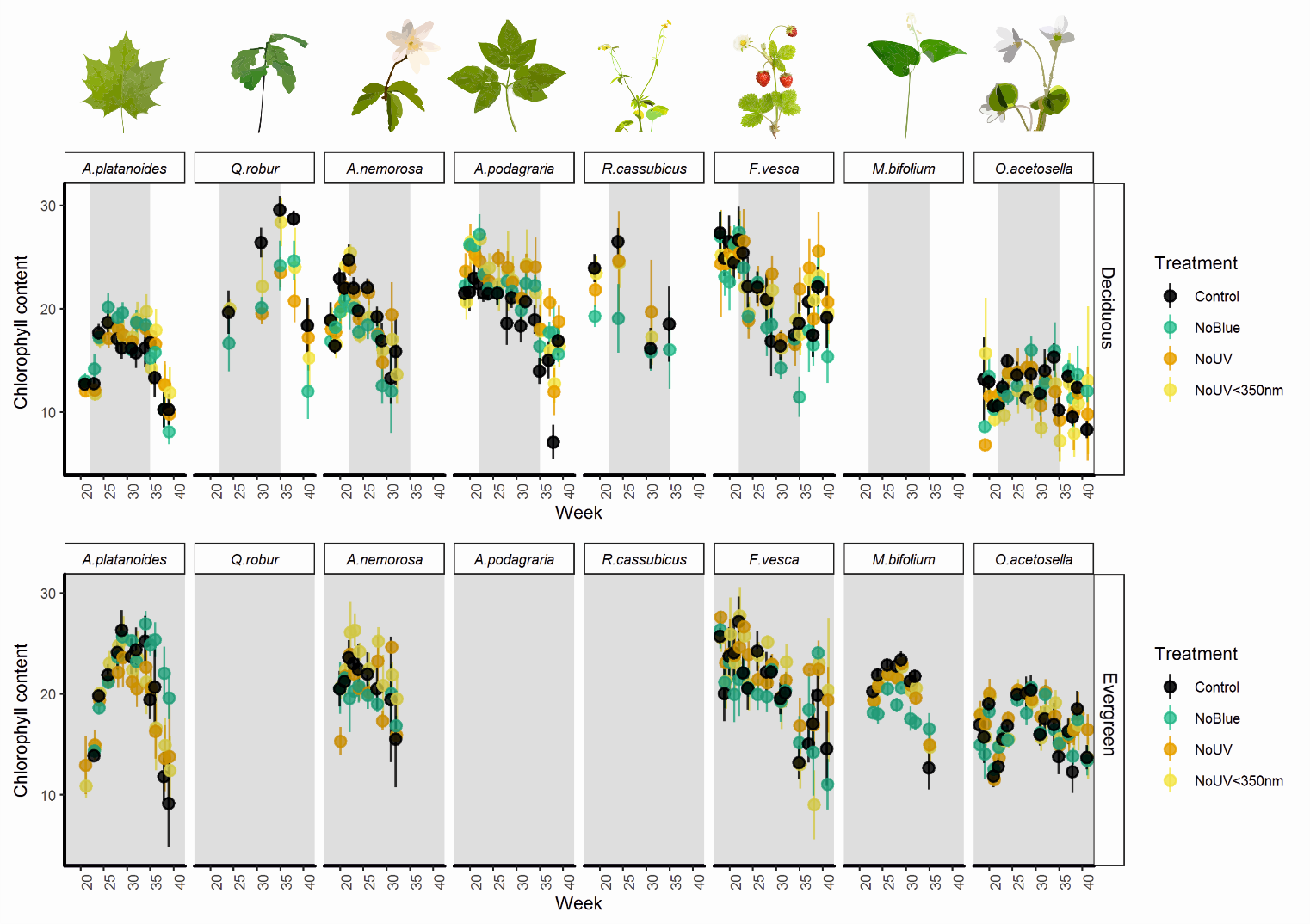

Figure S11. Chlorophyll content shown for eight different understorey plant species, growing during underneath filters in the understorey. The grey shaded area represents the time that canopy was closed during the measurement period. Means and 1±SE presented on the graph, with the individual plot as the unit of replication.

Table S3. ANOVA output from GAMM on *A. platanoides* leaf out.

| ***A. platanoides* leaf out** |  |  |  |
| --- | --- | --- | --- |
|  | df | F | p-value |
| StandType | 1 | 11.699 | 0.000653 |
| Treatment | 3 | 10.544 | 8.04E-07 |
| StandType:Treatment | 3 | 1.732 | 0.158839 |

Table S4. Summary output from GAMM on *A. platanoides* leaf out.

| ***A. platanoides* leaf out** |  |  |  |  |  |
| --- | --- | --- | --- | --- | --- |
|  | Estimate | Std. Error | t value | Pr(>\|t\|) |  |
| (Intercept) | 4.34737 | 0.07782 | 55.865 | < 2e-16 | *** |
| StandTypeEvergreen | -0.46859 | 0.137 | -3.42 | 0.000653 | *** |
| TreatmentNoBlue | -0.40546 | 0.08841 | -4.586 | 5.15E-06 | *** |
| TreatmentNoUV | 0.05505 | 0.08588 | 0.641 | 0.521693 |  |
| TreatmentNoUV<350nm | -0.15954 | 0.0866 | -1.842 | 0.065757 | . |
| StandTypeEvergreen:TreatmentNoBlue | 0.17632 | 0.15381 | 1.146 | 0.251948 |  |
| StandTypeEvergreen:TreatmentNoUV | -0.15013 | 0.15135 | -0.992 | 0.321489 |  |
| StandTypeEvergreen:TreatmentNoUV<350nm | 0.11243 | 0.1508 | 0.746 | 0.456118 |  |
| StandTypeEvergreen:TreatmentNoUV<350nm | 0.11243 | 0.1508 | 0.746 | 0.456118 |  |
|  | edf | Ref.df | F | p-value |  |
| s(Time) | 8.80E+00 | 8.797 | 657.42 | < 2e-16 | *** |
| ti(Time,Treatment) | 1.40E-06 | 12 | 0 | 0.603 |  |
| t2(YEAR,StandType,Stand,Time,Plot) | 1.11E+01 | 18 | 2.32 | 1.38E-06 | *** |
| **Comparison against UV filter** | Estimate | Std. Error | t value | Pr(>\|t\|) |  |
| (Intercept) | 4.39987 | 0.08064 | 54.565 | < 2e-16 | *** |
| StandTypeEvergreen | -0.62095 | 0.13632 | -4.555 | 5.94E-06 | *** |
| TreatmentControl | -0.05582 | 0.08605 | -0.649 | 0.5167 |  |
| TreatmentNoBlue | -0.46715 | 0.0899 | -5.196 | 2.50E-07 | *** |
| TreatmentNoUV<350nm | -0.21464 | 0.08792 | -2.441 | 0.0148 | * |
| StandTypeEvergreen:TreatmentControl | 0.15081 | 0.15166 | 0.994 | 0.3203 |  |
| StandTypeEvergreen:TreatmentNoBlue | 0.33327 | 0.15522 | 2.147 | 0.032 | * |
| StandTypeEvergreen:TreatmentNoUV<350nm | 0.2623 | 0.15209 | 1.725 | 0.0849 | . |

Table S5. ANOVA output from GAMM on *A. nemorosa* leaf out.

| ***A. nemorosa* leaf out** |  |  |  |
| --- | --- | --- | --- |
|  | df | F | p-value |
| StandType | 1 | 9.049 | 0.003 |
| Treatment | 3 | 2.434 | 0.0664 |
| StandType:Treatment | 3 | 0.219 | 0.8829 |

Table S6. Summary output from GAMM on *A. nemorosa* leaf out.

| ***A. nemorosa* leaf out** |  |  |  |  |  |
| --- | --- | --- | --- | --- | --- |
|  | Estimate | Std. Error | t value | Pr(>\|t\|) |  |
| (Intercept) | 1.88178 | 0.03861 | 48.732 | <2e-16 | *** |
| StandTypeEvergreen | -0.23393 | 0.07776 | -3.008 | 0.003 | ** |
| TreatmentNoBlue | -0.07353 | 0.05041 | -1.459 | 0.146 |  |
| TreatmentNoUV | -0.062 | 0.05277 | -1.175 | 0.242 |  |
| TreatmentNoUV<350nm | 0.05096 | 0.05155 | 0.989 | 0.324 |  |
| StandTypeEvergreen:TreatmentNoBlue | -0.06994 | 0.10801 | -0.648 | 0.518 |  |
| StandTypeEvergreen:TreatmentNoUV | 0.01555 | 0.10725 | 0.145 | 0.885 |  |
| StandTypeEvergreen:TreatmentNoUV<350nm | -0.01199 | 0.10447 | -0.115 | 0.909 |  |
|  | edf | Ref.df | F | p-value |  |
| s(Time) | 2.609 | 2.609 | 90.12 | <2e-16 | *** |
| t2(Time,StandType,Stand,Plot) | 4.007 | 13 | 0.58 | 0.0718 | . |

Table 7. ANOVA output from GAMM on *A. nemorosa* leaf out.

| ***A. podagraria* leaf out** |  |  |  |
| --- | --- | --- | --- |
|  | df | F | p-value |
| Treatment | 3 | 1.816 | 0.144 |

Table S8. ANOVA output from GAMM on *R. cassubicus* leaf out.

| ***R. cassubicus* leaf out** |  |  |  |
| --- | --- | --- | --- |
|  | df | F | p-value |
| Treatment | 3 | 0.924 | 0.43 |

Table S9. ANOVA output from GAMM on *A. platanoides* leaf senescence.

| ***A. platanoides* senescence** |  |  |  |
| --- | --- | --- | --- |
|  | df | F | p-value |
| StandType | 1 | 0.008 | 0.929763 |
| Treatment | 3 | 8.611 | 1.24E-05 |
| StandType:Treatment | 3 | 6.684 | 0.000184 |

Table S10. Summary output from GAMM on *A. platanoides* leaf senescence.

| ***A. platanoides* senescence** |  |  |  |  |  |
| --- | --- | --- | --- | --- | --- |
|  | Estimate | Std. Error | t value | Pr(>\|t\|) |  |
| (Intercept) | 2.95749 | 0.07938 | 37.258 | < 2e-16 | *** |
| StandTypeEvergreen | -0.01102 | 0.12497 | -0.088 | 0.92976 |  |
| TreatmentNoBlue | -0.40981 | 0.0825 | -4.968 | 8.20E-07 | *** |
| TreatmentNoUV | -0.26135 | 0.08214 | -3.182 | 0.00152 | ** |
| TreatmentNoUV<350nm | -0.26527 | 0.08167 | -3.248 | 0.00121 | ** |
| StandTypeEvergreen:TreatmentNoBlue | -0.35335 | 0.13373 | -2.642 | 0.00839 | ** |
| StandTypeEvergreen:TreatmentNoUV | 0.21916 | 0.12966 | 1.69 | 0.09133 | . |
| StandTypeEvergreen:TreatmentNoUV<350nm | 0.09923 | 0.12936 | 0.767 | 0.44325 |  |
|  | edf | Ref.df | F | p-value |  |
| s(Time) | 6.34 | 6.34 | 219.09 | < 2e-16 | *** |
| ti(Time,Treatment) | 7.213 | 12 | 3.703 | 2.09E-08 | *** |
| t2(YEAR,StandType,Stand,Time,Plot) | 10.199 | 15 | 3.581 | 5.47E-09 | *** |

| **Comparison against UV filter** |  |  |  |  |  |
| --- | --- | --- | --- | --- | --- |
|  | Estimate | Std. Error | t value | Pr(>\|t\|) |  |
| (Intercept) | 2.6967 | 0.08047 | 33.511 | < 2e-16 | *** |
| StandTypeEvergreen | 0.20584 | 0.12601 | 1.633 | 0.10274 |  |
| TreatmentControl | 0.26309 | 0.08424 | 3.123 | 0.00185 | ** |
| TreatmentNoBlue | -0.14413 | 0.08508 | -1.694 | 0.09062 | . |
| TreatmentNoUV<350nm | -0.00321 | 0.08424 | -0.038 | 0.96961 |  |
| StandTypeEvergreen:TreatmentControl | -0.21772 | 0.13297 | -1.637 | 0.10192 |  |
| StandTypeEvergreen:TreatmentNoBlue | -0.57265 | 0.13719 | -4.174 | 3.30E-05 | *** |
| StandTypeEvergreen:TreatmentNoUV<350nm | -0.12086 | 0.13274 | -0.911 | 0.36281 |  |

Table S11. ANOVA output from GAMM on *R. cassubicus* leaf senescence.

| ***R. cassubicus* senescence** |  |  |  |
| --- | --- | --- | --- |
|  | df | F | p-value |
| Treatment | 3 | 3.595 | 0.0164 |

Table S12. Summary output from GAMM on *R. cassubicus* leaf senescence

| ***R. cassubicus* senescence** |  |  |  |  |  |
| --- | --- | --- | --- | --- | --- |
|  | Estimate | Std. Error | t value | Pr(>\|t\|) |  |
| (Intercept) | 2.49253 | 0.08389 | 29.711 | < 2e-16 | *** |
| TreatmentNoBlue | -0.44614 | 0.1372 | -3.252 | 0.00158 | ** |
| TreatmentNoUV | -0.11893 | 0.13659 | -0.871 | 0.38605 |  |
| TreatmentNoUV<350nm | -0.18053 | 0.12055 | -1.498 | 0.1375 |  |
|  | edf | Ref.df | F | p-value |  |
| s(Time) | 3.31E+00 | 3.309 | 340.5 | <2e-16 | *** |
| t2(Time,Stand,Plot) | 2.44E-07 | 5 | 0 | 0.374 |  |
| **Comparison against UV filter** |  |  |  |  |  |
|  | Estimate | Std. Error | t value | Pr(>\|t\|) |  |
| (Intercept) | 2.3736 | 0.1078 | 22.015 | <2e-16 | *** |
| TreatmentControl | 0.1189 | 0.1366 | 0.871 | 0.386 |  |
| TreatmentNoBlue | -0.3272 | 0.1532 | -2.136 | 0.0352 | * |
| TreatmentNoUV<350nm | -0.0616 | 0.1382 | -0.446 | 0.6568 |  |

Table S13. ANOVA output from GAMM on *A. podagraria* leaf senescence.

| ***A. podagraria* senescence** |  |  |  |
| --- | --- | --- | --- |
|  | df | F | p-value |
| Treatment | 3 | 3.848 | 0.0114 |

Table S14. Summary output from GAMM on *A. podagraria* leaf senescence.

| ***A. podagraria* senescence** |  |  |  |  |  |
| --- | --- | --- | --- | --- | --- |
|  | Estimate | Std. Error | t value | Pr(>\|t\|) |  |
| (Intercept) | 2.65401 | 0.10321 | 25.714 | < 2e-16 | *** |
| TreatmentNoBlue | -0.33591 | 0.11478 | -2.927 | 0.00412 | ** |
| TreatmentNoUV | -0.27243 | 0.11177 | -2.437 | 0.01631 | * |
| TreatmentNoUV<350nm | -0.07927 | 0.11317 | -0.7 | 0.48507 |  |

|  | edf | Ref.df | F | p-value |  |
| --- | --- | --- | --- | --- | --- |
| s(Time) | 3.577 | 3.577 | 362.146 | <2e-16 | *** |
| t2(Time,Stand,Plot) | 3.421 | 5 | 4.628 | 8.00E-05 | *** |
| **Comparison against UV filter** |  |  |  |  |  |
|  | Estimate | Std. Error | t value | Pr(>\|t\|) |  |
| (Intercept) | 2.38158 | 0.11347 | 20.988 | <2e-16 | *** |
| TreatmentControl | 0.27243 | 0.11177 | 2.437 | 0.0163 | * |
| TreatmentNoBlue | -0.06348 | 0.11649 | -0.545 | 0.5869 |  |
| TreatmentNoUV<350nm | 0.19317 | 0.11469 | 1.684 | 0.0948 | . |

Table S15. ANOVA output from GAMM on *Q. robur* leaf senescence.

| ***Q. robur* senescence** |  |  |  |
| --- | --- | --- | --- |
|  | df | F | p-value |
| Treatment | 3 | 2.189 | 0.0949 |

| ***A. platanoides* adaxial flavonol** |  |  |  |  |  |
| --- | --- | --- | --- | --- | --- |
|  | Estimate | Std. Error | t value | Pr(>\|t\|) |  |
| (Intercept) | 0.9024 | 0.033401 | 27.017 | < 2e-16 | *** |
| StandTypeEvergreen | -0.45618 | 0.046954 | -9.716 | < 2e-16 | *** |
| TreatmentNoBlue | -0.43314 | 0.017849 | -24.266 | < 2e-16 | *** |
| TreatmentNoUV | -0.12648 | 0.017401 | -7.269 | 9.14E-13 | *** |
| TreatmentNoUV<350nm | -0.17045 | 0.017456 | -9.765 | < 2e-16 | *** |
| StandTypeEvergreen:TreatmentNoBlue | 0.248783 | 0.029508 | 8.431 | < 2e-16 | *** |
| StandTypeEvergreen:TreatmentNoUV | 0.004177 | 0.028484 | 0.147 | 0.8835 |  |
| StandTypeEvergreen:TreatmentNoUV<350nm | 0.046779 | 0.02811 | 1.664 | 0.0965 | . |
|  | edf | Ref.df | F | p-value |  |
| s(Week) | 8.183 | 8.183 | 5.36E+00 | 1.78E-06 |  |
| ti(Week,Treatment) | 12 | 12 | 0 | 1 |  |
| t2(YEAR,Week,StandType,Stand,PLOT) | 15.862 | 18 | 22.417 | < 2e-16 |  |
| **Comparison against UV filter** |  |  |  |  |  |
|  | Estimate | Std. Error | t value | Pr(>\|t\|) |  |
| (Intercept) | 0.775918 | 0.033454 | 23.194 | < 2e-16 | *** |
| StandTypeEvergreen | -0.452 | 0.047244 | -9.568 | < 2e-16 | *** |
| TreatmentControl | 0.126481 | 0.017401 | 7.269 | 9.14E-13 | *** |
| TreatmentNoBlue | -0.30665 | 0.017829 | -17.199 | < 2e-16 | *** |
| TreatmentNoUV<350nm | -0.04397 | 0.017476 | -2.516 | 0.0121 | * |
| StandTypeEvergreen:TreatmentControl | -0.00418 | 0.028484 | -0.147 | 0.8835 |  |
| StandTypeEvergreen:TreatmentNoBlue | 0.244607 | 0.029534 | 8.282 | 5.56E-16 | *** |
| StandTypeEvergreen:TreatmentNoUV<350nm | 0.042603 | 0.028386 | 1.501 | 0.1338 |  |

| ***A. platanoides* adaxial flavonols** |  |  |  |
| --- | --- | --- | --- |
|  | df | F | p-value |
| StandType | 1 | 94.39 | <2e-16 |
| Treatment | 3 | 205.66 | <2e-16 |
| StandType:Treatment | 3 | 30.9 | <2e-16 |

Table S16A. ANOVA output from GAMM on *A. platanoides* adaxial flavonols.

Table S16B. Summary output from GAMM on *A. platanoides* adaxial flavonols.

Table S16C. ANOVA output from GAMM on *A. podagraria* adaxial flavonols.

| ***A. podagraria* adaxial flavonols** |  |  |  |
| --- | --- | --- | --- |
|  | df | F | p-value |
| Treatment | 3 | 71.64 | <2e-16 |

Table S16D. Summary output from GAMM on *A. podagraria* adaxial flavonols.

| ***A. podagraria* adaxial flavonols** |  |  |  |  |  |
| --- | --- | --- | --- | --- | --- |
|  | Estimate | Std. Error | t value | Pr(>\|t\|) |  |
| (Intercept) | 0.74681 | 0.07024 | 10.632 | <2e-16 | *** |
| TreatmentNoBlue | -0.31161 | 0.02476 | -12.586 | <2e-16 | *** |
| TreatmentNoUV | -0.05332 | 0.02544 | -2.096 | 0.037 | * |
| TreatmentNoUV<350nm | -0.05287 | 0.02793 | -1.893 | 0.0594 | . |
|  | edf | Ref.df | F | p-value |  |
| s(Week):TreatmentControl | 5.126 | 5.126 | 6.62 | 9.54E-06 | *** |
| s(Week):TreatmentNoBlue | 1 | 1 | 1.4 | 0.237699 |  |
| s(Week):TreatmentNoUV | 3.507 | 3.507 | 4.07 | 0.003236 | ** |
| s(Week):TreatmentNoUV<350nm | 4.354 | 4.354 | 5.837 | 0.000174 | *** |
| ti(Week,Treatment) | 4.856 | 12 | 3.026 | 2.06E-08 | *** |
| t2(YEAR,Week,Stand,PLOT) | 6.82 | 9 | 12.015 | < 2e-16 | *** |
| **Comparison against UV filter** |  |  |  |  |  |
|  | Estimate | Std. Error | t value | Pr(>\|t\|) |  |
| (Intercept) | 0.720174 | 0.048602 | 14.818 | <2e-16 | *** |
| TreatmentControl | 0.046217 | 0.027805 | 1.662 | 0.0976 | . |
| TreatmentNoBlue | -0.26582 | 0.026666 | -9.968 | <2e-16 | *** |
| TreatmentNoUV<350nm | -0.00344 | 0.028985 | -0.119 | 0.9057 |  |

Table S16E. ANOVA output from LME on *A. nemorosa* adaxial flavonols.

| ***A. nemorosa* adaxial flavonols** |  |  |  |
| --- | --- | --- | --- |
| Parametric | Terms: |  |  |
|  | df | F | p-value |
| StandType | 1.00 | 209.90 | <2e-16 |
| Treatment | 3.00 | 252.26 | <2e-16 |
| StandType:Treatment | 3.00 | 35.15 | <2e-16 |

Table S16F. Summary output from LME on *A. nemorosa* adaxial flavonols.

| ***A. nemerosa* adaxial flavonols** |  |  |  |  |  |
| --- | --- | --- | --- | --- | --- |
|  | Estimate | Std. Error | t value | Pr(>\|t\|) |  |
| (Intercept) | 1.07667 | 0.02438 | 44.169 | < 2e-16 | *** |
| StandTypeEvergreen | -0.63778 | 0.04901 | -13.013 | < 2e-16 | *** |
| TreatmentNoBlue | -0.47333 | 0.01864 | -25.394 | < 2e-16 | *** |
| TreatmentNoUV | -0.13988 | 0.0181 | -7.73 | 6.72E-14 | *** |
| TreatmentNoUV<350nm | -0.09258 | 0.0177 | -5.232 | 2.54E-07 | *** |
| StandTypeEvergreen:TreatmentNoBlue | 0.36708 | 0.04124 | 8.902 | < 2e-16 | *** |
| StandTypeEvergreen:TreatmentNoUV | 0.04004 | 0.03708 | 1.08 | 0.2808 |  |
| StandTypeEvergreen:TreatmentNoUV<350nm | 0.06765 | 0.03735 | 1.811 | 0.0707 | . |
|  | edf | Ref.df | F | p-value |  |
| s(Week) | 7.264 | 7.264 | 20.328 | < 2e-16 | *** |
| ti(Week,Treatment) | 4.924 | 12 | 1.177 | 0.00482 | ** |
| t2(Week,StandType,Stand,REALPLOT) | 10.823 | 15 | 7.557 | < 2e-16 | *** |
| **Comparison against UV filter** |  |  |  |  |  |
|  | Estimate | Std. Error | t value | Pr(>\|t\|) |  |
| (Intercept) | 0.93678 | 0.02456 | 38.148 | < 2e-16 | *** |
| StandTypeEvergreen | -0.59774 | 0.04747 | -12.592 | < 2e-16 | *** |
| TreatmentControl | 0.13988 | 0.0181 | 7.73 | 6.72E-14 | *** |
| TreatmentNoBlue | -0.33345 | 0.01895 | -17.599 | < 2e-16 | *** |
| TreatmentNoUV<350nm | 0.0473 | 0.01814 | 2.608 | 0.00941 | ** |
| StandTypeEvergreen:TreatmentControl | -0.04004 | 0.03708 | -1.08 | 0.28079 |  |
| StandTypeEvergreen:TreatmentNoBlue | 0.32703 | 0.04063 | 8.049 | 7.08E-15 | *** |
| StandTypeEvergreen:TreatmentNoUV<350nm | 0.02761 | 0.03676 | 0.751 | 0.45303 |  |

Table S16G. ANOVA output from GAMM on *F. vesca* adaxial flavonols.

| ***F. vesca* adaxial flavonols** |  |  |  |
| --- | --- | --- | --- |
|  | df | F | p-value |
| StandType | 1 | 237.78 | < 2e-16 |
| Treatment | 3 | 12.51 | 5.42E-08 |
| StandType:Treatment | 3 | 1.04 | 0.374 |

Table S16H. Summary output from GAMM on *F. vesca* adaxial flavonols.

| ***F. vesca* adaxial flavonols** |  |  |  |  |  |
| --- | --- | --- | --- | --- | --- |
|  | Estimate | Std. Error | t value | Pr(>\|t\|) |  |
| (Intercept) | 0.924708 | 0.085554 | 10.808 | < 2e-16 | *** |
| StandTypeEvergreen | -0.95907 | 0.062196 | -15.42 | < 2e-16 | *** |
| TreatmentNoBlue | -0.16252 | 0.028204 | -5.762 | 1.20E-08 | *** |
| TreatmentNoUV | -0.13418 | 0.027362 | -4.904 | 1.14E-06 | *** |
| TreatmentNoUV<350nm | -0.08496 | 0.02806 | -3.028 | 0.00254 | ** |
| StandTypeEvergreen:TreatmentNoBlue | -0.00332 | 0.036372 | -0.091 | 0.9272 |  |
| StandTypeEvergreen:TreatmentNoUV | 0.045374 | 0.036063 | 1.258 | 0.20869 |  |
| StandTypeEvergreen:TreatmentNoUV<350nm | -0.00733 | 0.037981 | -0.193 | 0.84701 |  |
|  | edf | Ref.df | F | p-value |  |
| s(Week) | 6.981 | 6.981 | 32.414 | < 2e-16 | *** |
| ti(Week,Treatment) | 4.669 | 12 | 1.271 | 0.00134 | ** |
| t2(YEAR,Week,StandType,Stand,PLOT) | 18.693 | 21 | 11.717 | < 2e-16 | *** |
| **Comparison against UV filter** |  |  |  |  |  |
|  | Estimate | Std. Error | t value | Pr(>\|t\|) |  |
| (Intercept) | 0.955875 | 0.115894 | 8.248 | 6.87E-16 | *** |
| StandTypeEvergreen | -1.28688 | 0.071766 | -17.932 | < 2e-16 | *** |
| TreatmentControl | 0.100458 | 0.029989 | 3.35 | 0.000848 | *** |
| TreatmentNoBlue | -0.03501 | 0.03015 | -1.161 | 0.245897 |  |
| TreatmentNoUV<350nm | 0.027881 | 0.030545 | 0.913 | 0.361629 |  |
| StandTypeEvergreen:TreatmentControl | -0.00271 | 0.043483 | -0.062 | 0.950299 |  |
| StandTypeEvergreen:TreatmentNoBlue | -0.03284 | 0.043833 | -0.749 | 0.453994 |  |
| StandTypeEvergreen:TreatmentNoUV<350nm | -0.03105 | 0.045631 | -0.68 | 0.49644 |  |

Table S16I. ANOVA output from GAMM on *M. bifolium* adaxial flavonols.

| ***M. bifolium* adaxial flavonols** |  |  |  |
| --- | --- | --- | --- |
|  | df | F | p-value |
| Treatment | 3 | 23.22 | 1.42E-12 |

Table S16J. Summary output from GAMM on *M. bifolium* adaxial flavonols.

| ***M.bifolium* adaxial flavonols** |  |  |  |  |  |
| --- | --- | --- | --- | --- | --- |
|  | Estimate | Std. Error | t value | Pr(>\|t\|) |  |
| (Intercept) | 0.282182 | 0.005917 | 47.69 | < 2e-16 | *** |
| TreatmentNoBlue | -0.05069 | 0.006276 | -8.077 | 1.37E-13 | *** |
| TreatmentNoUV | -0.02296 | 0.005943 | -3.863 | 0.000161 | *** |
| TreatmentNoUV<350nm | -0.01718 | 0.006457 | -2.661 | 0.008573 | ** |
|  | edf | Ref.df | F | p-value |  |
| s(Week) | 3.683 | 3.683 | 17.886 | 7.73E-12 | *** |
| ti(Week,Treatment) | 2.875 | 12 | 0.449 | 0.133005 |  |
| t2(YEAR,Week,PLOT) | 3.683 | 6 | 3.234 | 0.000262 | *** |
| **Comparison against UV filter** |  |  |  |  |  |
|  | Estimate | Std. Error | t value | Pr(>\|t\|) |  |
| (Intercept) | 0.266396 | 0.005752 | 46.313 | < 2e-16 | *** |
| TreatmentControl | 0.041521 | 0.007565 | 5.488 | 1.51E-07 | *** |
| TreatmentNoBlue | -0.02177 | 0.008629 | -2.523 | 0.0126 | * |
| TreatmentNoUV<350nm | 0.004228 | 0.008288 | 0.51 | 0.6107 |  |

Table S16K. ANOVA output from GAMM on *O. acetosella* adaxial flavonols.

| ***O. acetosella* adaxial flavonols** |  |  |  |
| --- | --- | --- | --- |
|  | df | F | p-value |
| StandType | 1 | 175.25 | < 2e-16 |
| Treatment | 3 | 26 | 7.44E-16 |
| StandType:Treatment | 3 | 11.54 | 2.26E-07 |

Table S16L. Summary output from GAMM on *O. acetosella* adaxial flavonols.

| ***O. acetosella* adaxial flavonols** |  |  |  |  |  |
| --- | --- | --- | --- | --- | --- |
|  | Estimate | Std. Error | t value | Pr(>\|t\|) |  |
| (Intercept) | 0.73618 | 0.03249 | 22.659 | < 2e-16 | *** |
| StandTypeEvergreen | -0.36534 | 0.0276 | -13.238 | < 2e-16 | *** |
| TreatmentNoBlue | -0.12901 | 0.01548 | -8.336 | 4.92E-16 | *** |
| TreatmentNoUV | -0.03908 | 0.01529 | -2.555 | 0.010861 | * |
| TreatmentNoUV<350nm | -0.0216 | 0.01532 | -1.409 | 0.159254 |  |
| StandTypeEvergreen:TreatmentNoBlue | -0.02763 | 0.0178 | -1.552 | 0.121207 |  |
| StandTypeEvergreen:TreatmentNoUV | -0.09662 | 0.01767 | -5.469 | 6.56E-08 | *** |
| StandTypeEvergreen:TreatmentNoUV<350nm | -0.06723 | 0.01786 | -3.764 | 0.000183 | *** |
|  | edf | Ref.df | F | p-value |  |
| s(Week) | 7.462 | 7.462 | 48.121 | < 2e-16 | *** |
| ti(Week,Treatment) | 2.478 | 12 | 2.811 | 4.20E-08 | *** |
| t2(YEAR,Week,StandType,Stand,PLOT) | 11.94 | 14 | 9.771 | < 2e-16 | *** |
| **Comparison against UV filter** |  |  |  |  |  |
|  | Estimate | Std. Error | t value | Pr(>\|t\|) |  |
| (Intercept) | 0.70281 | 0.03244 | 21.667 | < 2e-16 | *** |
| StandTypeEvergreen | -0.4601 | 0.02686 | -17.133 | < 2e-16 | *** |
| TreatmentControl | 0.04176 | 0.01374 | 3.039 | 0.002471 | ** |
| TreatmentNoBlue | -0.08862 | 0.01506 | -5.883 | 6.56E-09 | *** |
| TreatmentNoUV<350nm | 0.01839 | 0.01408 | 1.306 | 0.191948 |  |
| StandTypeEvergreen:TreatmentControl | 0.10363 | 0.01684 | 6.156 | 1.34E-09 | *** |
| StandTypeEvergreen:TreatmentNoBlue | 0.06863 | 0.01796 | 3.822 | 0.000145 | *** |
| StandTypeEvergreen:TreatmentNoUV<350nm | 0.03288 | 0.01712 | 1.92 | 0.055294 | . |

Table S16M. ANOVA output from GAMM on *Q. robur* adaxial flavonols.

| ***Q. robur* adaxial flavonols** | Terms: |  |  |
| --- | --- | --- | --- |
|  | df | F | p-value |
| Treatment | 3 | 19.89 | 1.04E-09 |

Table S16N. Summary output from GAMM on *Q. robur* adaxial flavonols.

| ***Q. robur* adaxial flavonols** |  |  |  |  |  |
| --- | --- | --- | --- | --- | --- |
|  | Estimate | Std.Error | t-value | p-value |  |
| (Intercept) | 0.61733 | 0.03405 | 18.132 | < 2e-16 | *** |
| TreatmentNoBlue | -0.27244 | 0.03577 | -7.616 | 3.33E-11 | *** |
| TreatmentNoUV | -0.2455 | 0.03661 | -6.706 | 2.08E-09 | *** |
| TreatmentNoUV>350nm | -0.17692 | 0.04001 | -4.422 | 2.88E-05 | *** |
| --- |  |  |  |  |  |
|  | edf | Ref.df | F | p-value |  |
| s(Week):TreatmentControl | 1 | 1 | 20.653 | 1.71E-05 | *** |
| s(Week):TreatmentNoBlue | 1 | 1 | 7.487 | 0.00753 | ** |
| s(Week):TreatmentNoUV | 1 | 1 | 2.639 | 0.10794 |  |
| s(Week):TreatmentNoUV>350nm | 1 | 1 | 2.792 | 0.09838 | . |
| t2(Week,Stand,REALPLOT) | 4.384 | 7 | 4.364 | 3.29E-05 | *** |
| **Comparison against UV filter** |  |  |  |  |  |
|  | Estimate | Std. Error | t value | Pr(>\|t\|) |  |
| (Intercept) | 0.37334 | 0.03986 | 9.367 | 5.20E-14 | *** |
| TreatmentControl | 0.24197 | 0.039 | 6.205 | 3.31E-08 | *** |
| TreatmentNoBlue | -0.02019 | 0.03882 | -0.52 | 0.6045 |  |
| TreatmentNoUV<350nm | 0.0737 | 0.04226 | 1.744 | 0.0855 | . |

Table S16O. ANOVA output from GAMM on *R. cassubicus* adaxial flavonols.

| ***R. cassubicus* adaxial flavonols** | df | F | p-value | p-value |
| --- | --- | --- | --- | --- |
| Treatment | 3 | 11.98 | 5.80E-06 | <.0001 |

Table S16P. Summary output from GAMM on *R. cassubicus* adaxial flavonols.

| ***R. cassubicus* adaxial flavonols** |  |  |  |  |  |
| --- | --- | --- | --- | --- | --- |
|  | Value | Std.Error | DF | t-value | p-value |
| (Intercept) | 0.8375563 | 0.138761 | 54 | 6.035975 | 0 |
| Week | -0.0055535 | 0.005304 | 54 | -1.04695 | 0.2998 |
| TreatmentNoBlue | -0.5109987 | 0.213684 | 54 | -2.39138 | 0.0203 |
| TreatmentNoUV | -0.0583252 | 0.215603 | 54 | -0.27052 | 0.7878 |
| TreatmentNoUV>350nm | -0.0201342 | 0.177674 | 54 | -0.11332 | 0.9102 |
| Week:TreatmentNoBlue | 0.0075988 | 0.008037 | 54 | 0.945491 | 0.3486 |
| Week:TreatmentNoUV | -0.0038703 | 0.008831 | 54 | -0.43826 | 0.6629 |
| Week:TreatmentNoUV>350nm | -0.0036133 | 0.00688 | 54 | -0.52521 | 0.6016 |
|  | edf | Ref.df | F | p-value |  |
| s(Week) | 2.031 | 2.031 | 3.511 | 0.04482 | * |
| t2(Week,REALPLOT) | 3.191 | 5 | 3.163 | 0.00363 | ** |
| **Comparison against UV filter** |  |  |  |  |  |
|  | Estimate | Std. Error | t value | Pr(>\|t\|) |  |
| (Intercept) | 0.56925 | 0.05334 | 10.671 | 4.01E-14 | *** |
| TreatmentControl | 0.15212 | 0.0557 | 2.731 | 0.00887 | ** |
| TreatmentNoBlue | -0.17831 | 0.06192 | -2.879 | 0.00599 | ** |
| TreatmentNoUV<350nm | 0.04168 | 0.0546 | 0.763 | 0.44906 |  |

Table S17A. ANOVA output from GAMM on *A. platanoides* adaxial anthocyanins.

| ***A. platanoides* adaxial anthocyanins** |  |  |  |
| --- | --- | --- | --- |
|  | df | F | p-value |
| StandType | 1 | 16.718 | 4.79E-05 |
| Treatment | 3 | 8.673 | 1.15E-05 |
| StandType:Treatment | 3 | 1.076 | 0.359 |

Table S17B. Summary output from GAMM on *A. platanoides* adaxial anthocyanins.

| ***A. platanoides* adaxial Anth** |  |  |  |  |  |
| --- | --- | --- | --- | --- | --- |
|  | Estimate | Std. Error | t value | Pr(>\|t\|) |  |
| (Intercept) | 0.430039 | 0.007289 | 58.999 | < 2e-16 | *** |
| StandTypeEvergreen | -0.04735 | 0.011581 | -4.089 | 4.79E-05 | *** |
| TreatmentNoBlue | -0.03395 | 0.007017 | -4.837 | 1.59E-06 | *** |
| TreatmentNoUV | -0.01472 | 0.006924 | -2.126 | 0.03379 | * |
| TreatmentNoUV<350nm | -0.02498 | 0.00691 | -3.615 | 0.00032 | *** |
| StandTypeEvergreen:TreatmentNoBlue | -0.00451 | 0.0111 | -0.406 | 0.68487 |  |
| StandTypeEvergreen:TreatmentNoUV | 0.009915 | 0.010938 | 0.906 | 0.36497 |  |
| StandTypeEvergreen:TreatmentNoUV<350nm | 0.012416 | 0.010714 | 1.159 | 0.24687 |  |
|  | edf | Ref.df | F | p-value |  |
| s(Week) | 6.8118 | 6.812 | 63.804 | < 2e-16 | *** |
| ti(Week,Treatment) | 0.9165 | 12 | 0.122 | 0.208 |  |
| t2(YEAR,Week,StandType,Stand,PLOT) | 13.066 | 18 | 5.161 | 3.81E-15 | *** |
| **Comparison against UV filter** |  |  |  |  |  |
|  | Estimate | Std. Error | t value | Pr(>\|t\|) |  |
| (Intercept) | 0.181391 | 0.006903 | 26.277 | <2e-16 | *** |
| StandTypeEvergreen | -0.02368 | 0.011361 | -2.084 | 0.0375 | * |
| TreatmentControl | 0.017295 | 0.006802 | 2.543 | 0.0112 | * |
| TreatmentNoBlue | -0.01669 | 0.006997 | -2.386 | 0.0173 | * |
| TreatmentNoUV<350nm | -0.00398 | 0.006851 | -0.581 | 0.5613 |  |
| StandTypeEvergreen:TreatmentControl | -0.00589 | 0.011109 | -0.53 | 0.596 |  |
| StandTypeEvergreen:TreatmentNoBlue | -0.02153 | 0.011538 | -1.866 | 0.0624 | . |
| StandTypeEvergreen:TreatmentNoUV<350nm | -0.00579 | 0.011074 | -0.523 | 0.6013 |  |

Table S17C. ANOVA output from GAMM on *A. podagraria* adaxial anthocyanins.

| ***A. podagraria* adaxial anthocyanins** |  |  |  |
| --- | --- | --- | --- |
|  | df | F | p-value |
| Treatment | 3 | 6.657 | 0.000231 |

Table S17D. Summary output from GAMM on *A. podagraria* adaxial anthocyanins.

| ***A. podagraria* adaxial anthocyanins** |  |  |  |  |  |
| --- | --- | --- | --- | --- | --- |
|  | Estimate | Std. Error | t value | Pr(>\|t\|) |  |
| (Intercept) | 0.149643 | 0.005707 | 26.222 | < 2e-16 | *** |
| TreatmentNoBlue | -0.01985 | 0.005118 | -3.878 | 0.00013 | *** |
| TreatmentNoUV | -0.02248 | 0.00545 | -4.125 | 4.84E-05 | *** |
| TreatmentNoUV<350nm | -0.01769 | 0.005967 | -2.965 | 0.00328 | ** |
|  | edf | Ref.df | F | p-value |  |
| s(Week):TreatmentControl | 3.22E+00 | 3.219 | 33.322 | < 2e-16 | *** |
| s(Week):TreatmentNoBlue | 2.19E+00 | 2.186 | 26.503 | 8.37E-12 | *** |
| s(Week):TreatmentNoUV | 2.99E+00 | 2.985 | 22.524 | 3.12E-13 | *** |
| s(Week):TreatmentNoUV<350nm | 3.13E+00 | 3.132 | 23.822 | 2.51E-14 | *** |
| ti(Week,Treatment) | 1.13E-07 | 12 | 0 | 0.0767 | . |
| t2(YEAR,Week,Stand,PLOT) | 4.42E+00 | 9 | 3.427 | 2.56E-06 | *** |
| **Comparison against UV filter** |  |  |  |  |  |
|  | Estimate | Std. Error | t value | Pr(>\|t\|) |  |
| (Intercept) | 0.130152 | 0.005546 | 23.469 | < 2e-16 | *** |
| TreatmentControl | 0.020653 | 0.005167 | 3.997 | 8.10E-05 | *** |
| TreatmentNoBlue | 0.002684 | 0.004968 | 0.54 | 0.589 |  |
| TreatmentNoUV<350nm | 0.005544 | 0.00539 | 1.029 | 0.305 |  |

Table S17E. ANOVA output from LME on *A. nemorosa* adaxial anthocyanins.

| ***A. nemorosa* adaxial anthocyanins** |  |  |  |  |
| --- | --- | --- | --- | --- |
|  | numDF | denDF | F-value | p-value |
| (Intercept) | 1 | 40 | 1231.783 | <.0001 |
| Week | 1 | 40 | 6.2043 | 0.017 |
| StandType | 1 | 14 | 2.7508 | 0.1194 |
| Treatment | 3 | 40 | 0.5344 | 0.6613 |
| StandType:Treatment | 3 | 40 | 1.1978 | 0.3229 |

Table S17F. Summary output from LME on *A. nemorosa* adaxial anthocyanins.

| ***A. nemorosa* adaxial anthocyanins** |  |  |  |  |  |
| --- | --- | --- | --- | --- | --- |
|  | Value | Std.Error | DF | t-value | p-value |
| (Intercept) | 0.3670226 | 0.077436 | 40 | 4.739662 | 0 |
| Week | -0.0067697 | 0.003307 | 40 | -2.04714 | 0.0473 |
| StandTypeEvergreen | -0.0480043 | 0.024198 | 14 | -1.98378 | 0.0672 |
| TreatmentNoBlue | -0.0325862 | 0.017299 | 40 | -1.88367 | 0.0669 |
| TreatmentNoUV | -0.0177469 | 0.017433 | 40 | -1.01801 | 0.3148 |
| TreatmentNoUV>350nm | -0.0106209 | 0.01731 | 40 | -0.61358 | 0.543 |
| StandTypeEvergreen:TreatmentNoBlue | 0.0611929 | 0.03682 | 40 | 1.661934 | 0.1043 |
| StandTypeEvergreen:TreatmentNoUV | 0.0397424 | 0.032296 | 40 | 1.23057 | 0.2257 |
| StandTypeEvergreen:TreatmentNoUV>350nm | 0.0103153 | 0.033809 | 40 | 0.305103 | 0.7619 |

Table S17G. ANOVA output from GAMM on *F. vesca* adaxial anthocyanins.

| ***F. vesca* adaxial anthocyanins** |  |  |  |
| --- | --- | --- | --- |
|  | df | F | p-value |
| StandType | 1 | 8.115 | 0.00451 |
| Treatment | 3 | 3.769 | 0.01052 |

Table S17H. Summary output from GAMM on *F. vesca* adaxial anthocyanins.

| ***F. vesca* adaxial anthocyanins** |  |  |  |  |  |
| --- | --- | --- | --- | --- | --- |
|  | Estimate | Std. Error | t value | Pr(>\|t\|) |  |
| (Intercept) | 0.1402 | 0.006683 | 20.979 | < 2e-16 | *** |
| StandTypeEvergreen | -0.02609 | 0.00916 | -2.849 | 0.00451 | ** |
| TreatmentNoBlue | 0.008418 | 0.005289 | 1.592 | 0.11187 |  |
| TreatmentNoUV | -0.00512 | 0.005122 | -0.999 | 0.3182 |  |
| TreatmentNoUV<350nm | -0.00718 | 0.005255 | -1.365 | 0.17252 |  |
| StandTypeEvergreen:TreatmentNoBlue | -0.00557 | 0.006898 | -0.808 | 0.41944 |  |
| StandTypeEvergreen:TreatmentNoUV | -0.00777 | 0.006853 | -1.134 | 0.25708 |  |
| StandTypeEvergreen:TreatmentNoUV<350nm | 0.002933 | 0.007188 | 0.408 | 0.68334 |  |
|  | edf | Ref.df | F | p-value |  |
| s(Week) | 6.42E+00 | 6.421 | 15.982 | <2e-16 | *** |
| ti(Week,Treatment) | 2.57E-08 | 12 | 0 | 0.478 |  |
| t2(YEAR,Week,StandType,Stand,PLOT) | 1.54E+01 | 21 | 9.494 | <2e-16 | *** |

Table S17I. ANOVA output from GAMM on *M. bifolium* adaxial anthocyanins.

| ***M. bifolium* adaxial anthocyanins** |  |  |  |
| --- | --- | --- | --- |
|  | df | F | p-value |
| Treatment | 3 | 3.933 | 0.00966 |

Table S17J. Summary output from GAMM on *M. bifolium* adaxial anthocyanins.

| ***M.bifolium* adaxial anthocyanins** |  |  |  |  |  |
| --- | --- | --- | --- | --- | --- |
|  | Estimate | Std. Error | t value | Pr(>\|t\|) |  |
| (Intercept) | 0.124143 | 0.003862 | 32.148 | <2e-16 | *** |
| TreatmentNoBlue | 0.014348 | 0.0049 | 2.928 | 0.0039 | ** |
| TreatmentNoUV | 0.000102 | 0.004222 | 0.024 | 0.9807 |  |
| TreatmentNoUV<350nm | -0.00139 | 0.004761 | -0.292 | 0.7709 |  |
|  | edf | Ref.df | F | p-value |  |
| s(Week) | 3.787 | 3.787 | 25.502 | < 2e-16 | *** |
| ti(Week,Treatment) | 6.006 | 12 | 1.659 | 0.0028 | ** |
| t2(YEAR,Week,PLOT) | 3.158 | 6 | 2.022 | 0.00386 | ** |
| **Comparison against UV filter** |  |  |  |  |  |
|  | Estimate | Std. Error | t value | Pr(>\|t\|) |  |
| (Intercept) | 1.24E-01 | 3.50E-03 | 35.491 | < 2e-16 | *** |
| TreatmentControl | 1.92E-05 | 4.17E-03 | 0.005 | 0.99633 |  |
| TreatmentNoBlue | 1.45E-02 | 4.77E-03 | 3.046 | 0.00271 | ** |
| TreatmentNoUV<350nm | -1.43E-03 | 4.58E-03 | -0.313 | 0.75491 |  |

Table S17K. ANOVA output from GAMM on *O. acetosella* adaxial anthocyanins.

| ***O. acetosella* adaxial anthocyanins** |  |  |  |
| --- | --- | --- | --- |
|  | df | F | p-value |
| StandType | 1 | 17.507 | 3.27E-05 |
| Treatment | 3 | 5.241 | 0.00141 |
| StandType:Treatment | 3 | 6.519 | 0.00024 |

Table S17L. Summary output from GAMM on *O. acetosella* adaxial anthocyanins.

| ***O. acetosella* adaxial anthocyanins** |  |  |  |  |  |
| --- | --- | --- | --- | --- | --- |
|  | Estimate | Std. Error | t value | Pr(>\|t\|) |  |
| (Intercept) | 0.177683 | 0.006398 | 27.773 | < 2e-16 | *** |
| StandTypeEvergreen | -0.03201 | 0.007651 | -4.184 | 3.27E-05 | *** |
| TreatmentNoBlue | -0.00417 | 0.006731 | -0.619 | 0.5363 |  |
| TreatmentNoUV | 0.0127 | 0.007024 | 1.808 | 0.07109 | . |
| TreatmentNoUV<350nm | 0.020448 | 0.00707 | 2.892 | 0.00396 | ** |
| StandTypeEvergreen:TreatmentNoBlue | 0.007269 | 0.007424 | 0.979 | 0.32793 |  |
| StandTypeEvergreen:TreatmentNoUV | -0.01711 | 0.007718 | -2.217 | 0.02697 | * |
| StandTypeEvergreen:TreatmentNoUV<350nm | -0.02107 | 0.007825 | -2.693 | 0.00727 | ** |
|  | edf | Ref.df | F | p-value |  |
| s(Week) | 7.52E+00 | 7.517 | 35.172 | < 2e-16 | *** |
| ti(Week,Treatment) | 8.65E-08 | 12 | 0 | 0.992 |  |
| t2(YEAR,Week,StandType,Stand,PLOT) | 7.70E+00 | 14 | 5.483 | 5.41E-14 | *** |
| **Comparison against UV filter** |  |  |  |  |  |
|  | Estimate | Std. Error | t value | Pr(>\|t\|) |  |
| (Intercept) | 0.191035 | 0.006614 | 28.884 | < 2e-16 | *** |
| StandTypeEvergreen | -0.05005 | 0.007993 | -6.261 | 7.08E-10 | *** |
| TreatmentControl | -0.01289 | 0.005578 | -2.31 | 0.021219 | * |
| TreatmentNoBlue | -0.01842 | 0.006161 | -2.99 | 0.002902 | ** |
| TreatmentNoUV<350nm | 0.003228 | 0.005797 | 0.557 | 0.577896 |  |
| StandTypeEvergreen:TreatmentControl | 0.019365 | 0.00687 | 2.819 | 0.004974 | ** |
| StandTypeEvergreen:TreatmentNoBlue | 0.025853 | 0.007362 | 3.512 | 0.000477 | *** |
| StandTypeEvergreen:TreatmentNoUV<350nm | 0.001967 | 0.007053 | 0.279 | 0.780363 |  |

Table S17M. ANOVA output from GAMM on *Q. robur* adaxial anthocyanins.

| ***Q. robur* adaxial anthocyanins** |  |  |  |
| --- | --- | --- | --- |
| Parametric | Terms: |  |  |
|  | df | F | p-value |
| Treatment | 3 | 3.351 | 0.023 |

Table S17N. Summary output from GAMM on *Q. robur* adaxial anthocyanins.

| ***Q. robur* adaxial anthocyanins** |  |  |  |  |  |
| --- | --- | --- | --- | --- | --- |
|  | Estimate | Std.Error | t-value | p-value |  |
| (Intercept) | 0.36354 | 0.01245 | 29.198 | <2e-16 | *** |
| TreatmentNoBlue | 0.03902 | 0.01772 | 2.203 | 0.0306 | * |
| TreatmentNoUV | 0.01667 | 0.01485 | 1.123 | 0.265 |  |
| TreatmentNoUV>350nm | 0.0299 | 0.01875 | 1.595 | 0.1148 |  |
| --- |  |  |  |  |  |
|  | edf | Ref.df | F | p-value |  |
| s(Week) | 1 | 1 | 7.743 | 0.00687 | ** |
| ti(Week,Treatment) | 9 | 9 | 0 | 1 |  |
| t2(Week,Stand,PLOT) | 4.384 | 7 | 4.364 | 3.29E-05 | *** |
| **Comparison against UV filter** |  |  |  |  |  |
|  | Estimate | Std. Error | t value | Pr(>\|t\|) |  |
| (Intercept) | 0.165914 | 0.013298 | 12.477 | <2e-16 | *** |
| TreatmentControl | -0.01613 | 0.018534 | -0.871 | 0.387 |  |
| TreatmentNoBlue | 0.016364 | 0.018088 | 0.905 | 0.369 |  |
| TreatmentNoUV<350nm | 0.006054 | 0.019846 | 0.305 | 0.761 |  |

Table S17O. ANOVA output from GAMM on *R. cassubicus* adaxial anthocyanins.

| ***R. cassubicus* adaxial anthocyanins** |  |  |  |  |
| --- | --- | --- | --- | --- |
|  | df | F | p-value | p-value |
| Treatment | 3 | 0.097 | 0.962 | <.0001 |

Table S17P. Summary output from GAMM on *R. cassubicus* adaxial anthocyanins.

| ***R. cassubicus* adaxial anthocyanins** |  |  |  |  |  |
| --- | --- | --- | --- | --- | --- |
|  | Value | Std.Error | DF | t-value | p-value |
| (Intercept) | 0.02805407 | 0.031568 | 54 | 0.888684 | 0.3781 |
| Week | 0.00420982 | 0.001205 | 54 | 3.493703 | 0.001 |
| TreatmentNoBlue | 0.01123649 | 0.048385 | 54 | 0.232229 | 0.8172 |
| TreatmentNoUV | 0.04951649 | 0.048836 | 54 | 1.013931 | 0.3151 |
| TreatmentNoUV>350nm | -0.00934154 | 0.040202 | 54 | -0.23237 | 0.8171 |
| Week:TreatmentNoBlue | -0.0004974 | 0.001821 | 54 | -0.27317 | 0.7858 |
| Week:TreatmentNoUV | -0.00234265 | 0.002 | 54 | -1.17117 | 0.2467 |
| Week:TreatmentNoUV>350nm | 0.00028692 | 0.001557 | 54 | 0.184315 | 0.8545 |
|  | edf | Ref.df | F | p-value |  |
| s(Week) | 2.791 | 2.791 | 13.912 | 1.29E-05 | *** |
| t2(Week,REALPLOT) | 3.407 | 5 | 3.704 | 0.00199 | ** |
| **Comparison against UV filter** |  |  |  |  |  |
|  | Estimate | Std. Error | t value | Pr(>\|t\|) |  |
| (Intercept) | 0.133915 | 0.006173 | 21.694 | < 2e-16 | *** |
| StandTypeEvergreen | -0.03206 | 0.008606 | -3.725 | 0.000209 | *** |
| TreatmentControl | 0.00814 | 0.004718 | 1.725 | 0.084873 | . |
| TreatmentNoBlue | 0.014653 | 0.004703 | 3.116 | 0.001903 | ** |
| TreatmentNoUV<350nm | 0.003572 | 0.004791 | 0.746 | 0.456139 |  |
| StandTypeEvergreen:TreatmentControl | 0.006999 | 0.006864 | 1.02 | 0.308218 |  |
| StandTypeEvergreen:TreatmentNoBlue | 0.001023 | 0.006893 | 0.148 | 0.882061 |  |
| StandTypeEvergreen:TreatmentNoUV<350nm | 0.004039 | 0.00719 | 0.562 | 0.574386 |  |
